# Identification and correction of phase switches with Hi-C data in the Nanopore and HiFi chromosome-scale assemblies of the dikaryotic leaf rust fungus *Puccinia triticina*

**DOI:** 10.1101/2021.04.28.441890

**Authors:** Hongyu Duan, Ashley W. Jones, Tim Hewitt, Amy Mackenzie, Yiheng Hu, Anna Sharp, David Lewis, Rohit Mago, Narayana M. Upadhyaya, John P. Rathjen, Eric A. Stone, Benjamin Schwessinger, Melania Figueroa, Peter N. Dodds, Sambasivam Periyannan, Jana Sperschneider

**Affiliations:** Biological Data Science Institute, The Australian National University, Canberra, Australia; Research School of Biology, The Australian National University, Canberra, Australia; Black Mountain Science and Innovation Park, CSIRO Agriculture and Food, Canberra, Australia; John Curtin School of Medical Research, The Australian National University, Canberra, Australia

**Keywords:** Long-read sequencing, Hi-C, chromosomes, phasing, phase switches, HiFi, Nanopore, genome assembly

## Abstract

**Background:** Most animals and plants have more than one set of chromosomes and package these haplotypes into a single nucleus within each cell. In contrast, many fungal species carry multiple haploid nuclei per cell. Rust fungi are such species with two nuclei (karyons) that contain a full set of haploid chromosomes each. The physical separation of haplotypes in dikaryons means that, unlike in diploids, Hi-C chromatin contacts between haplotypes are false positive signals.

**Results:** We generate the first chromosome-scale, fully-phased assembly for the dikaryotic leaf rust fungus *Puccinia triticina* and compare Nanopore MinION and PacBio HiFi sequence-based assemblies. We show that false positive Hi-C contacts between haplotypes are predominantly caused by phase switches rather than by collapsed regions or Hi-C read mis-mappings. We introduce a method for phasing of dikaryotic genomes into the two haplotypes using Hi-C contact graphs, including a phase switch correction step. In the HiFi assembly, relatively few phase switches occur, and these are predominantly located at haplotig boundaries and can be readily corrected. In contrast, phase switches are widespread throughout the Nanopore assembly. We show that haploid genome read coverage of 30-40 times using HiFi sequencing is required for phasing of the leaf rust genome (~0.7% heterozygosity) and that HiFi sequencing resolves genomic regions with low heterozygosity that are otherwise collapsed in the Nanopore assembly.

**Conclusions:** This first Hi-C based phasing pipeline for dikaryons and comparison of long-read sequencing technologies will inform future genome assembly and haplotype phasing projects in other non-haploid organisms.

## Background

Genome assemblies that are as close as possible to the biological truth are the foundation for high-quality downstream functional studies and comparative analyses at scale. Recent advances in long-read genome sequencing technologies, such as Pacific Biosciences (PacBio) and Oxford Nanopore Technologies (ONT), have improved quality of genome assemblies by allowing the capture of more sequence information than previous commonly used approaches (Amarasinghe *et al.*, 2020). The ONT MinION is a portable, real-time DNA and RNA sequencing device that delivers long reads lengths of 10–100 Kb or even longer (Lang *et al.*, 2020). The Nanopore sequencing device outputs an electrical current signal which is translated to sequencing reads by basecalling software. However, basecalled reads currently have high error rates of ~5-20% (Chen *et al.*, 2021). In contrast, PacBio High-Fidelity (HiFi) sequencing, which outputs shorter read lengths (~10-20 Kb) than ONT, provides accuracy as high as Illumina short reads (> 99.9%). In combination with scaffolding data (Hi-C, optical maps or genetic linkage maps) chromosome-scale assemblies are now achievable for many species from whole genome long-read sequencing data (Logsdon *et al.*, 2020; Michael & VanBuren, 2020; Zhang *et al.*, 2020). However, phasing of haplotypes within a heterozygous diploid genome remains challenging. Current scaffolding methods applied to unphased, non-haploid assemblies lead to false positive fusions of allelic contigs (Zhang *et al.*, 2019). Noisy Hi-C signals between allelic contigs in the haplotypes can also be caused by mis-assemblies, collapsed regions, phase switches and difficulties of mapping in homologous or repetitive regions of the genome. Importantly, in the absence of a highly accurate phased reference genomes or parental data it can be difficult to quantify the rate of these errors in newly generated genome assemblies.

Whilst animals and plants package their diploid and polyploid genomes into a single nucleus, rust fungi, like many other fungi, contain two distinct haploid nuclei (dikaryons) with no physical contact between the homologous chromosomes (Lorrain *et al.*, 2019). The physical separation of haplotypes in dikaryons makes these systems ideal for assessing mis-assemblies and phase switch errors in non-haploid genome assemblies using Hi-C chromatin contact information. One of these dikaryotic rust fungi is *Puccinia triticina* (*Pt*), the causative agent of leaf rust. Leaf rust is one of the most damaging and widely distributed diseases of wheat worldwide (Figueroa *et al.*, 2017). It is caused by a macrocyclic, heteroecious, dikaryotic rust fungus with five spore stages (Bolton *et al.*, 2008; Lorrain *et al.*, 2019). During the asexual phase of *Pt* on the wheat host, urediniospores are deposited on the leaf surface by wind or rain and germinate. Appressoria form and penetration occurs through stomata with subsequent development of specialized infection structures called haustoria, that enable nutrient uptake as well as the delivery of effector proteins into the host plant cell (Garnica *et al.*, 2014). At approximately 7-10 days post infection (dpi), urediniospores are produced and erupt through the leaf surface to reinitiate the infection cycle. *Pt* can cycle indefinitely as uredinial infections on its wheat host as long as environmental conditions are favourable (Bolton *et al.*, 2008). The alternate host of leaf rust, *Thalictrum*, is rarely present in wheat-growing areas worldwide, so sexual recombination is unlikely a significant contributor to genetic variation in leaf rust (Kolmer, 2005). From a biological perspective, understanding the factors underpinning genome evolution in *Pt* has captured the interest of the scientific community. However, chromosome-scale, fully-phased assemblies for this species are thus far not available preventing to address these research questions.

The haploid genome sizes of rust fungi range from ~80Mb to ~2Gb (Tavares *et al.*, 2014; Ramos *et al.*, 2015; Figueroa *et al.*, 2020). Repetitive regions and the presence of two homologous haplotypes in these organisms often lead to assembly errors. Thus, rust genome assemblies from short reads have common limitations of being highly fragmented and being an underestimation of the true genome size. For example, two *Pt* short-read assemblies of races 77 and 106 have more than 44,000 contigs and assembled genome sizes of only ~100 Mb (Kiran *et al.*, 2016) and the American *Pt* isolate 1-1 BBBD Race 1 was assembled into 135.4 Mb with 21.3% gaps using a Roche 454 and Sanger sequencing strategy (N/L50: 68/544.256 Kb) (Cuomo *et al.*, 2017). The first PacBio long-read assembly for *Pt* (Australian isolate *Pt*104) achieved a 140.5 Mb primary assembly (N/L50: 23/2.073 Mb) with 128 Mb of associated haplotigs; however it presented a high percentage of duplicated single-copy ortholog genes (~12%) in the primary assembly suggesting the haplotypes were not fully resolved (Wu *et al.*, 2020).

Long-read data alone is insufficient for achieving chromosome-scale assemblies in rust fungi. Chromatin contact data such as Hi-C is essential for phasing of the two haplotypes and achieving chromosome-scale scaffolding. Across all rust fungi only the genome of the stem rust fungus *Puccinia graminis* f. sp. *tritici* (*Pgt*) has thus far been fully phased into the two haplotype chromosomes sets. This assembly resulted from a combination of PacBio RSII long read sequence for assembly, parental data available from a natural hybridization event involving a single nucleus exchange between isolates for nuclear haplotype assignment, and Hi-C data for scaffolding (Li, F. *et al.*, 2019). A fully-phased, chromosome-scale assembly of *Pt* or any other rust fungus is not available. Furthermore, full phasing of a rust fungus genome assembly using Hi-C data alone has not been performed thus far.

## Results

### HiFi technology resolves the two leaf rust haplotypes whilst Nanopore collapses ~12% of the assembly

An Australian isolate of the leaf rust fungus (pathotype 76-3,5,7,9,10,12,13) was collected from wheat cultivar *Morocco*. This isolate, hereafter referred to as *Pt76*, was used for genome assembly using two distinct approaches: firstly with Nanopore long-read sequencing in combination with Illumina short-read sequences for polishing, and secondly with HiFi long-read sequencing (Table 1). For the Nanopore sequenced based assembly we obtained a total of 6.7 Gb of Nanopore reads (L50 of reads: 30.7 Kb), which were assembled using Canu (Koren *et al.*, 2017), and subsequently polished and cleaned. This process yielded an assembly of 717 contigs with total size of 233.2 Mb and N/L50 of 35/1.1 Mb, with 95.7% of BUSCOs present. The estimated *k*-mer completeness (fraction of reliable *k*-mers in the Illumina read set that are also found in the assembly) determined by merqury (Rhie *et al.*, 2020) is 95.8%. We also generated a separate assembly from 10.8 Gb HiFi reads (L50 of reads: 15.1 Kb) with either the Canu or hifiasm assemblers (Cheng *et al.*, 2021). After cleaning, the HiFi-Canu assembly contained 600 contigs with total size of 256.5 Mb and N/L50 of 26/2.4 Mb, with 96.2% of BUSCOs present. The estimated *k*-mer completeness is 96.4%, thus slightly higher than that of the Nanopore assembly. The HiFi-hifiasm assembly yielded a very similar output to the HiFi-Canu assembly, with 608 contigs at total size of 260.3 Mb and N/L50 of 24/3.0 Mb, with 96.4% of BUSCOs present and an estimated *k*-mer completeness of 96.4%. Despite their shorter average lengths, the high accuracy and higher coverage of the HiFi reads allowed for an assembly of longer contigs than the Nanopore reads.

**Table 1:**
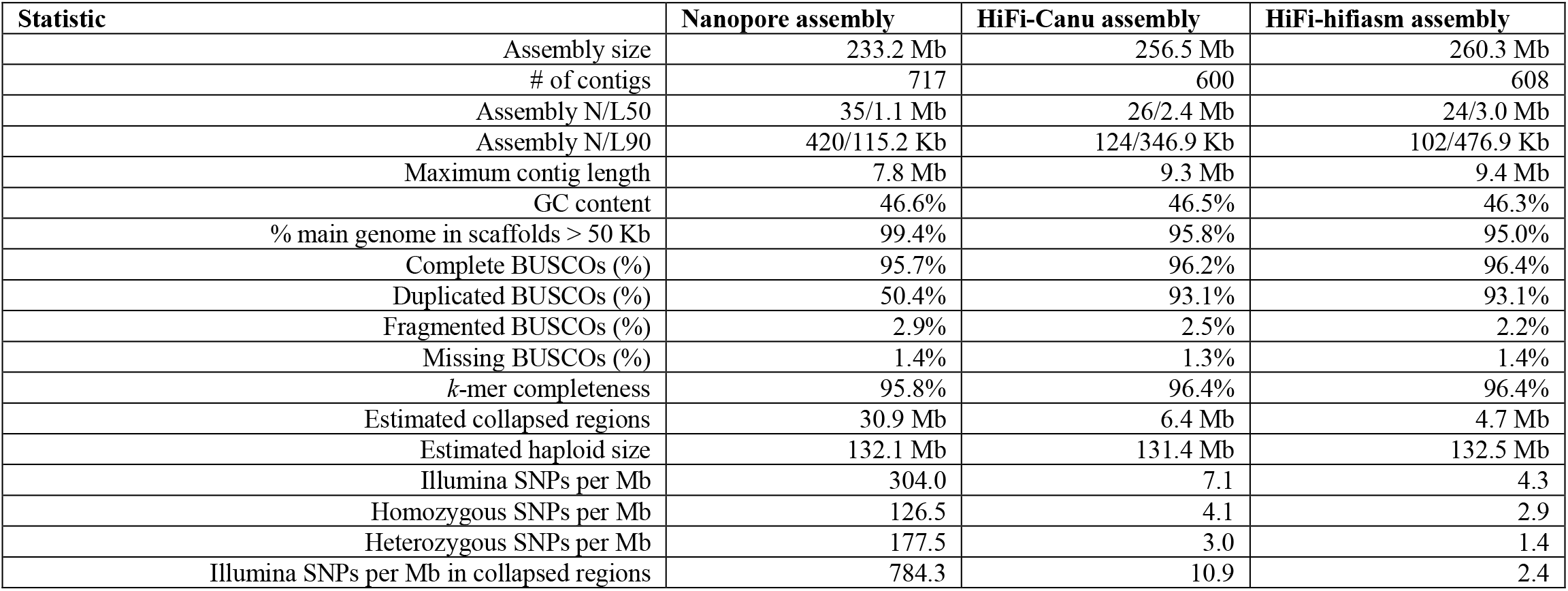
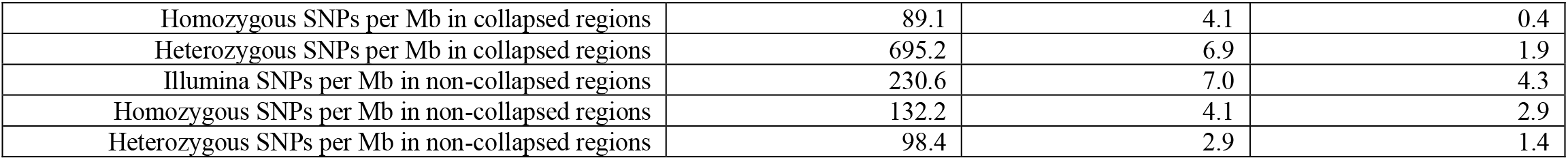
Statistics for the Nanopore and HiFi genome assemblies of *Pt76*. Statistics are shown for the clean assemblies with mitochondrial contigs, low-coverage contigs and contaminants removed. Assembly sizes and BUSCO statistics indicate that the two haplotypes are resolved in the HiFi assemblies. SNP calling shows that the HiFi assemblies are a highly accurate representation of the haplotypes, whereas the Nanopore assembly is a pseudo-haplotype representation of the genome.

| Statistic | Nanopore assembly | HiFi-Canu assembly | HiFi-hifiasm assembly |
| --- | --- | --- | --- |
| Assembly size | 233.2 Mb | 256.5 Mb | 260.3 Mb |
| # of contigs | 717 | 600 | 608 |
| Assembly N/L50 | 35/1.1 Mb | 26/2.4 Mb | 24/3.0 Mb |
| Assembly N/L90 | 420/115.2 Kb | 124/346.9 Kb | 102/476.9 Kb |
| Maximum contig length | 7.8 Mb | 9.3 Mb | 9.4 Mb |
| GC content | 46.6% | 46.5% | 46.3% |
| % main genome in scaffolds > 50 Kb | 99.4% | 95.8% | 95.0% |
| Complete BUSCOs (%) | 95.7% | 96.2% | 96.4% |
| Duplicated BUSCOs (%) | 50.4% | 93.1% | 93.1% |
| Fragmented BUSCOs (%) | 2.9% | 2.5% | 2.2% |
| Missing BUSCOs (%) | 1.4% | 1.3% | 1.4% |
| *k*-mer completeness | 95.8% | 96.4% | 96.4% |
| Estimated collapsed regions | 30.9 Mb | 6.4 Mb | 4.7 Mb |
| Estimated haploid size | 132.1 Mb | 131.4 Mb | 132.5 Mb |
| Illumina SNPs per Mb | 304.0 | 7.1 | 4.3 |
| Homozygous SNPs per Mb | 126.5 | 4.1 | 2.9 |
| Heterozygous SNPs per Mb | 177.5 | 3.0 | 1.4 |
| Illumina SNPs per Mb in collapsed regions | 784.3 | 10.9 | 2.4 |
| Homozygous SNPs per Mb in collapsed regions | 89.1 | 4.1 | 0.4 |
| Heterozygous SNPs per Mb in collapsed regions | 695.2 | 6.9 | 1.9 |
| Illumina SNPs per Mb in non-collapsed regions | 230.6 | 7.0 | 4.3 |
| Homozygous SNPs per Mb in non-collapsed regions | 132.2 | 4.1 | 2.9 |
| Heterozygous SNPs per Mb in non-collapsed regions | 98.4 | 2.9 | 1.4 |

We investigated the efficiency and accuracy of generating haplotypes assemblies using each sequencing technology (Nanopore vs. HiFi long reads). A long-read coverage analysis estimated that in the Nanopore assembly ~30.9 Mb are collapsed genomic regions (Figure 1A), representing about 12% of the assembly. In contrast, the HiFi-Canu and HiFi-hifiasm assemblies only had an estimated 6.4 Mb (Figure 1B) and 4.7 Mb collapsed genomic regions (Figure 1C) respectively. The HiFi assemblies are thus the closest complete representation of the two haplotypes. This is supported by the substantially higher proportion of duplicated BUSCOs in the HiFi assemblies at 93% compared to 50% in the Nanopore assembly (Table 1). Based on these collapsed regions, the estimated haploid genome size in each case is about 132 Mb. This is consistent with estimates from GenomeScope used with short-read Illumina data (Vurture *et al.*, 2017) which also indicates that the haploid genome size could be ~132 Mb.

**Figure 1:**
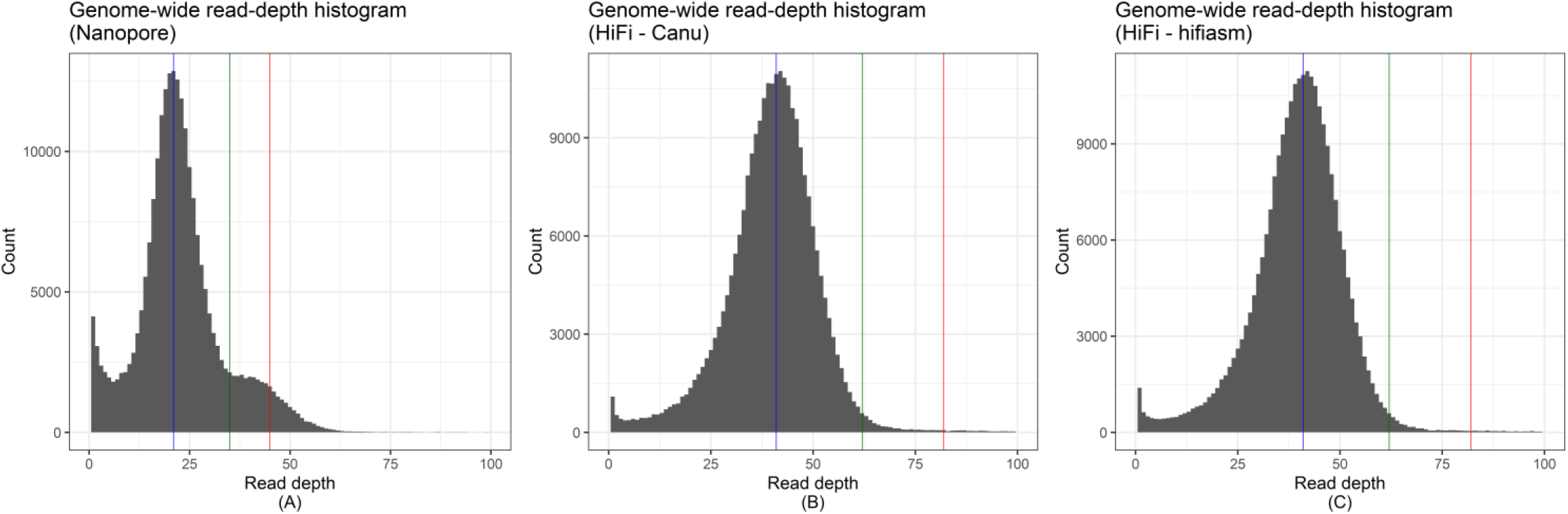
Genome-wide read-depth histograms of the (A) Nanopore and (B) HiFi assemblies. An assembly with collapsed regions will show a bimodal distribution with a first peak at haploid coverage (indicated by blue line) and a second peak at 2x haploid coverage (indicated by red line). The green line represents the mid-point and genomic regions with coverage above the mid-point are here defined as collapsed regions (Nanopore: 35x coverage; HiFi: 62x coverage). The HiFi assemblies appear to be fully phased whereas the Nanopore assembly has collapsed regions.

We used Illumina read mapping and SNP calling against all genome assemblies to assess the accuracy of each assembly. The HiFi assemblies have high accuracy with only ~6 SNPs per Mb (HiFi-Canu: 7.1 SNPs per Mb; HiFi-hifiasm: 4.3 SNPs per Mb), compared to the Nanopore assembly with ~300 SNPs per Mb. As expected, heterozygous SNPs are enriched in collapsed regions (Nanopore: 695.2 SNPs per Mb; HiFi-Canu: 6.9 SNPs per Mb; HiFi-hifiasm: 1.9 SNPs per Mb). Homozygous SNPs in non-collapsed regions indicate assembly accuracy, and in these regions the Nanopore assembly has 132 SNPs per Mb compared to 4 SNPs per Mb for the HiFi-Canu assembly and 3 SNPs per Mb for the HiFi-hifiasm assembly. These results indicate that the HiFi assemblies are a highly accurate representation of the entire genome of *Pt76*.

Lastly, we used Hi-C chromatin contact data to detect contig mis-joins in the Nanopore and HiFi assemblies and allow breaking of such chimeric contigs as part of genome reference curation steps and prevent erroneous scaffolding. In the Nanopore assembly, visual inspection of Hi-C contact maps for contigs >= 1Mb identified five mis-assemblies. We determined breakpoints based on lack of contiguity in the Nanopore long-read alignments to the contigs. For example, tig00000001 (11.9 Mb) is a chimeric contig with two centromeric regions visible in the Hi-C contact map (Supplementary Figure S1). In the HiFi-Canu assembly, we observed only one mis-joined contig from the Hi-C contact maps. We did not observe obvious chimeric contigs in the HiFi-hifiasm assembly.

### The HiFi assemblies have significantly less phase switches than the Nanopore assembly

In most plant and animal genomes, haplotypes reside in the same nucleus and thus Hi-C signals between homologous chromosomes are expected. However, in the dikaryotic rust fungi, haplotypes are physically separated in two nuclei which leads to no Hi-C signal between them. Thus, Hi-C signal between haplotypes in dikaryons will be a result of assembly errors, Hi-C read mis-mappings, collapsed regions and phase switch errors. To assess the rate of false positive Hi-C signal in the Nanopore and HiFi assemblies, we first constructed a highly confident subset of the two haplotypes that are expected to reside in separate nuclei. For this, we developed a gene binning method to find sets of homologous contigs which represent the two haplotypes (Figure 2). Genes that map exactly twice to the unphased assembly were used as phasing markers to assign homologous contigs into diploid scaffold bins *Bin_1_,…,Bin_n_*. Scaffold bins were constructed with a graph network approach where nodes are contigs and edges are the number of shared genes per Mb. Each strongly connected community in the graph is a diploid scaffold bin *Bin_x_* and contains two subsets *Bin_x_a_* and *Bin_x_b_*. Thus, a scaffold bin is part of a chromosome where the two subsets represent the two haplotypes.

**Figure 2:**
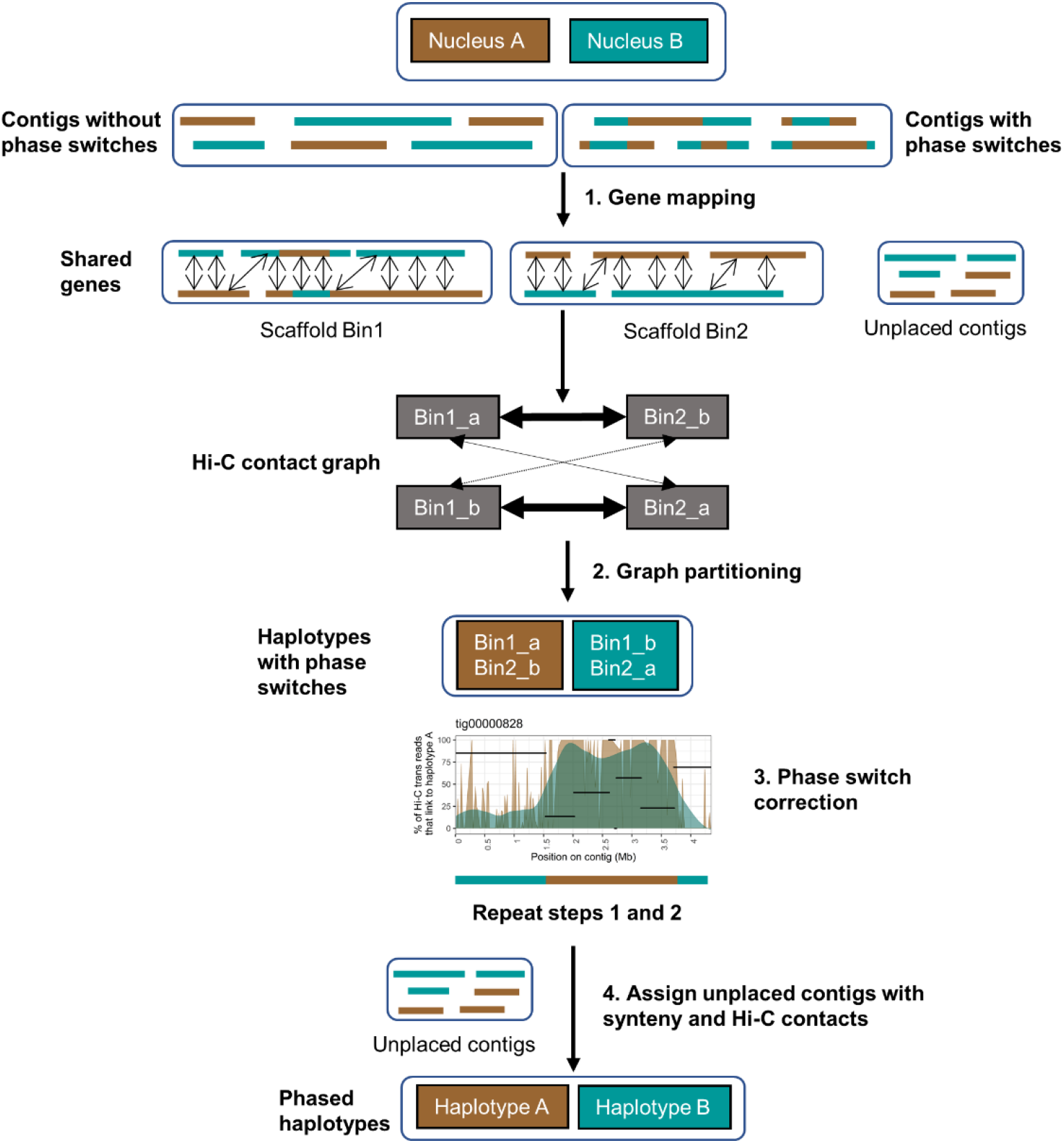
The dikaryotic phasing pipeline for assigning contigs into the two haplotypes A and B. A gene binning method was developed to phase the assembly into the two haplotypes prior to scaffolding of each haplotype separately. Genes that map exactly twice to the unphased assembly were used to assign contigs into scaffold bins that represent the two haplotypes. Hi-C contacts within allelic scaffold bins were ignored and the bins were clustered into the two haplotypes with a graph approach. Phase switches within scaffold bins were visualized, manually corrected and steps 1 and 2 were run again. Finally, unplaced contigs are assigned to the haplotypes based on synteny and Hi-C contacts.

As a test case, we first applied the gene binning method to the stem rust (*Pgt* 21-0) polished CANU assembly (PacBio RSII, 410 contigs, 176.9 Mb) (Li, F. *et al.*, 2019). The haploid genomes of *Pgt* 21-0 are highly heterozygous at ~15 SNPs/Kb and the CANU assembly has 31 contigs with 37 phase switches that were broken during subsequent fully-phased chromosome curation (Li, F. *et al.*, 2019). The *Pgt* assembly with these phase switches present produced 47 scaffold bins (149.1 Mb of the assembly, 27.8 Mb remain unassigned) while the *Pgt* assembly with the phase switch contigs broken (445 contigs, 176.9 Mb) produced 54 scaffold bins (145.7 Mb of the assembly, 31.2 Mb remain unassigned). Both *Pgt* assemblies were used in the following as controls in the Hi-C based phasing procedure. For leaf rust, 39 scaffold bins (202.5 Mb of the assembly, 30.8 Mb remain unassigned) were generated for the Nanopore assembly. In contrast, the HiFi-Canu assembly produced 26 scaffold bins (240.5 Mb of the assembly, 16.0 Mb remain unassigned) and the HiFi-hifiasm assembly produced 23 scaffold bins (242.0 Mb of the assembly, 18.2 Mb remain unassigned).

We used the scaffold binning to assess the false positive Hi-C contacts between haplotypes. We recorded normalized Hi-C contact frequencies for two sets: 1) *trans*-contacts between haplotypes (between contig subsets in a scaffold bin e.g. *Bin_1_a_* and *Bin_1_b_*, Figure 2); and 2) all other *trans*-contacts (e.g. between *Bin_1_a_* and *Bin_2_b_*). The proportion of *trans*-contacts that occur between haplotypes captures the false positive Hi-C signal between chromosomes in different nuclei. First, we investigated the distribution of mapping qualities for the Hi-C reads mapped between haplotypes. Mapping quality (MAPQ) reflects the degree of confidence in the point of origin for a read. For example, MAPQ of 10 or less indicates that there is at least a 1 in 10 chance that the read originated from another genomic location. In the *Pgt* and *Pt76* HiFi assemblies, the false positive Hi-C read mappings (*trans*-contacts between haplotypes) have lower mapping qualities than all other *trans*-contacts (*Pgt* mean: 17; HiFi-Canu mean: 14; HiFi-hifiasm mean: 14), with a large proportion of reads with MAPQ 0-10 (Figure 3). The opposite trend is observed in the *Pt76* Nanopore assembly, where the Hi-C read mappings for *trans*-contacts between haplotypes have high MAPQ with a peak around 30 (mean: 24). This shows that filtering Hi-C reads based on mapping quality reduces noisy Hi-C contacts between haplotypes, but only with a limited effect for the Nanopore assembly.

**Figure 3:**
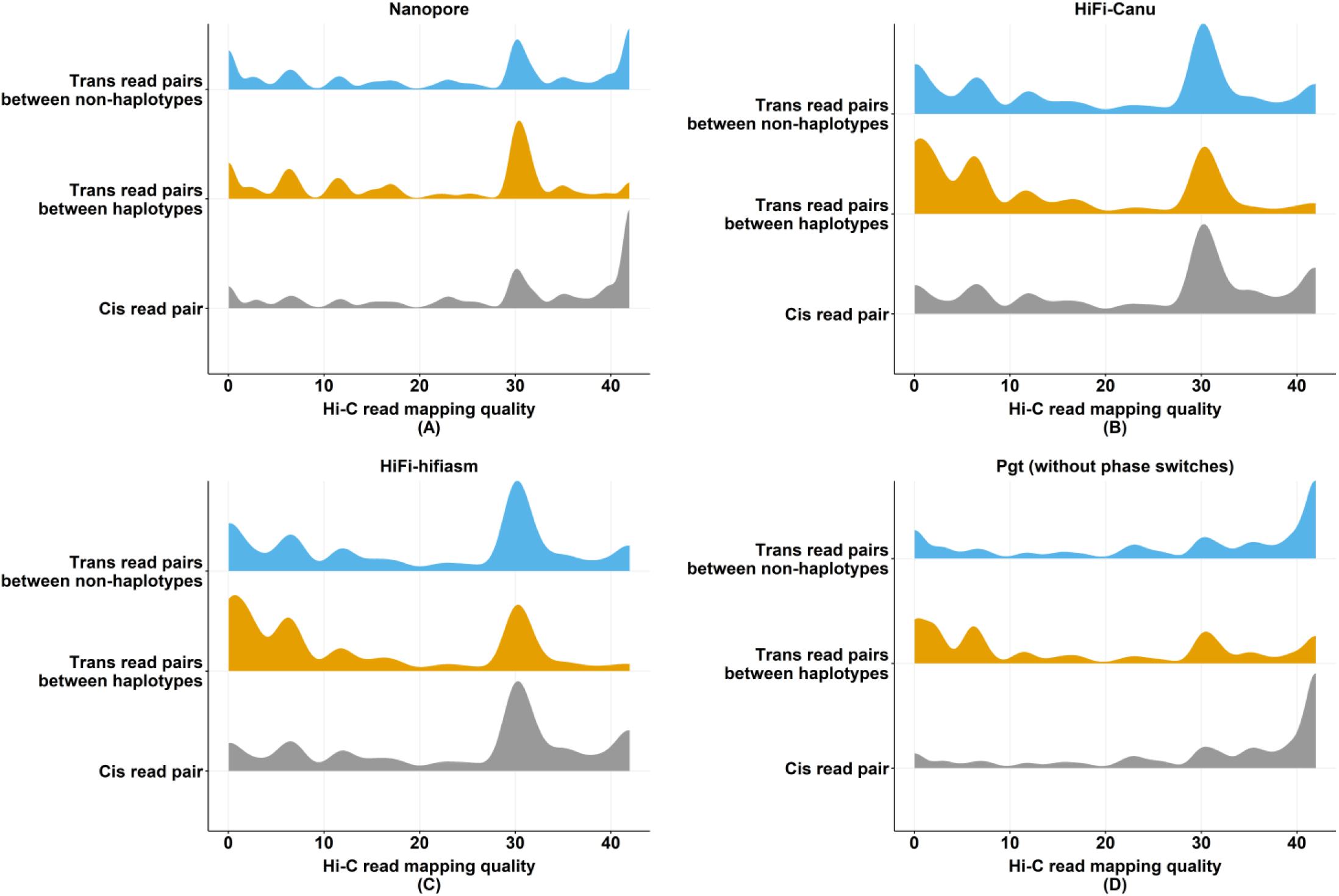
Distribution of mapping qualities for Hi-C reads. Density plots are shown for the distribution of mapping qualities for Hi-C read mappings with HiC-Pro (bowtie2, maximum possible MAPQ 42). In the **(A)** Nanopore assembly, the majority of *trans*-contacts between haplotypes have high MAPQ around 30, whereas they have lower MAPQ of 0-10 in the **(B)** and **(C)** HiFi assemblies.

For example, MAPQ of 10 or less indicates that there is at least a 1 in 10 chance that the read originated from another genomic location. In the *Pgt* assembly, the false positive Hi-C read mappings (*trans*-contacts between haplotypes) have lower mapping qualities (mean 17) than all other *trans*-contacts or *cis*-contacts, with a large proportion of reads with MAPQ <10 (Figure 3A). This suggests that most of the false positive contacts in this haplotype-resolved assembly result from poorly mapped reads and that filtering for read mapping quality above 30 should remove most of this background. A similar trend is observed in the *Pt76* Hifi assemblies, with a higher proportion of low quality mapping between haplotypes compared to all other *trans*-contacts or *cis*-contacts (Figure 3B,C). However, the *Pt76* Nanopore assembly showed a similar proportion of low quality read mappings for links between haplotypes and between bins, with both higher than for *cis*-contacts (Figure 3D), providing a first indication that the haplotypes in this assembly may be poorly resolved. In the following analysis of phase switch errors we only consider Hi-C read mappings where both reads have MAPQ of at least 30.

As a benchmark, we first assessed the rate of false positive *trans*-contacts between haplotypes in the *Pgt* assembly. In the raw assembly containing phase switch contigs, 6.4% of *trans*-contacts were between contigs of different haplotype assignment. In the corrected assembly after breaking the chimeric contigs, only 2.7% of *trans*-contacts were between haplotypes. The HiFi-Canu and HiFi-hifiasm *Pt76* assemblies gave similar results here to the *Pgt* uncorrected assembly with 8.6% and 7.3% of *trans*-contacts between opposite haplotypes (Table 2). This suggests that they may contain relatively few contigs with phase switches. However, in the *Pt76* Nanopore assembly we found that a high proportion (58.8%) of *trans*-contacts occur between haplotypes. This indicates that either extensive phase swapping occurs in the Nanopore assembly or that the large collapsed regions cause this false positive Hi-C signal. In the unpolished Nanopore assembly, the proportion of *trans*-contacts between haplotypes was slightly lower at 52.3%, indicating that while polishing might have introduced some local phase switch errors, this cannot explain most of the cross-haplotype contacts. Counterintuitively, we found that more Hi-C reads mapped in the Nanopore assembly than in the HiFi assemblies (Table 2). This can be explained by the handling of multi-mapped Hi-C reads. The HiFi assemblies where both haplotypes including homologous regions are represented will have higher rates of multi-mapped reads. Our mapping pipeline does not allow multi-mapped Hi-C reads, which leads to an exclusion of those reads where the haplotypes share near-identical sequence and thus overall lower Hi-C read mappings in the HiFi assemblies.

**Table 2:**
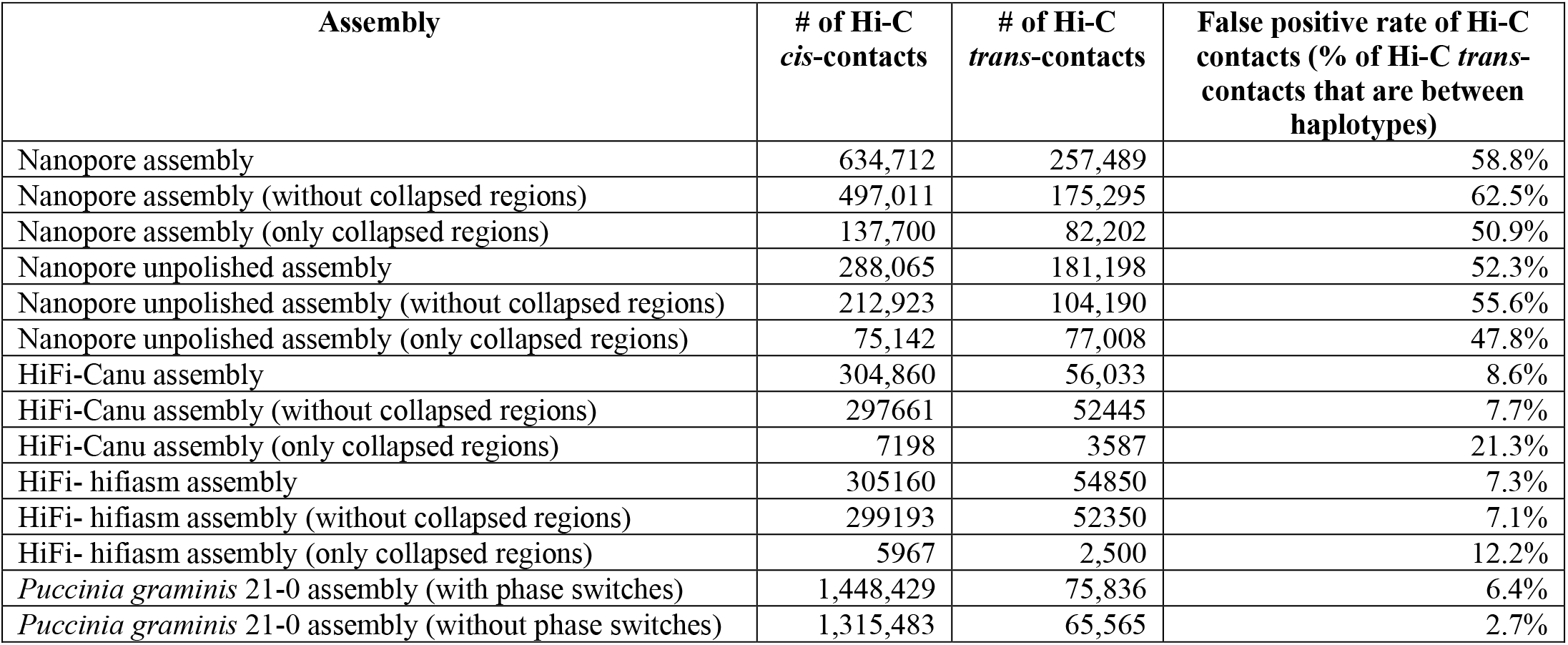
Proportion of false positive Hi-C contacts between haplotypes in the assemblies. Only Hi-C reads with high mapping quality (MAPQ > 30) were used in this analysis. Over half of all *trans* Hi-C contacts in the Nanopore assembly are false positive signals between haplotypes. In contrast, the HiFi assemblies have a low false positive signal similarly to the *Pgt* assembly with phase switches present.

| Assembly | # of Hi-C *cis*-contacts | # of Hi-C *trans*-contacts | False positive rate of Hi-C contacts (% of Hi-C *trans*-contacts that are between haplotypes) |
| --- | --- | --- | --- |
| Nanopore assembly | 634,712 | 257,489 | 58.8% |
| Nanopore assembly (without collapsed regions) | 497,011 | 175,295 | 62.5% |
| Nanopore assembly (only collapsed regions) | 137,700 | 82,202 | 50.9% |
| Nanopore unpolished assembly | 288,065 | 181,198 | 52.3% |
| Nanopore unpolished assembly (without collapsed regions) | 212,923 | 104,190 | 55.6% |
| Nanopore unpolished assembly (only collapsed regions) | 75,142 | 77,008 | 47.8% |
| HiFi-Canu assembly | 304,860 | 56,033 | 8.6% |
| HiFi-Canu assembly (without collapsed regions) | 297,661 | 52,445 | 7.7% |
| HiFi-Canu assembly (only collapsed regions) | 7,198 | 3,587 | 21.3% |
| HiFi-hifiasm assembly | 305,160 | 54,850 | 7.3% |
| HiFi-hifiasm assembly (without collapsed regions) | 299,193 | 52,350 | 7.1% |
| HiFi-hifiasm assembly (only collapsed regions) | 5,967 | 2,500 | 12.2% |
| *Puccinia graminis* 21-0 assembly (with phase switches) | 1,448,429 | 75,836 | 6.4% |
| *Puccinia graminis* 21-0 assembly (without phase switches) | 1,315,483 | 65,565 | 2.7% |

In collapsed regions and their surroundings, false positive Hi-C contacts between haplotypes are expected because both haplotypes are represented by a single sequence which has been placed into one haplotype but is absent from the other haplotype. We therefore excluded collapsed regions from the Hi-C contact analysis and expected that this should lead to less *trans*-contacts between haplotypes. For the HiFi-Canu and HiFi-hifiasm assemblies, excluding collapsed regions only slightly decreased the proportion of *trans*-contacts that occur between haplotypes to 7.7% and 7.1%, respectively (Table 2). Unexpectedly, for the Nanopore assembly excluding collapsed regions increased the proportion of *trans*-contacts between haplotypes to 62.5%. Thus, whilst collapsed regions may contribute to the noisy phasing signal in the Nanopore assembly, this analysis strongly suggests that there are extensive phase switches in the Nanopore assembly.

### Phase switches occur in large blocks and mostly at haplotig boundaries in the HiFi assemblies

We developed an algorithm to separate the binned contigs into two haplotype sets representing their nuclear origin. To do this, we constructed a graph based on Hi-C links between the scaffold bins, ignoring Hi-C links within scaffold bins (Figure 2). A graph network approach then returned the two expected communities that represent a high proportion of the phased haplotypes, which might still include phase switches. We first validated the phasing of the scaffold bins using the phase-switch corrected *Pgt* assembly as the nuclear origin of each contig was previously determined based on parental sequence data from two isolates involved in a nuclear exchange event (Li et al 2019). The Hi-C links between the two haplotypes of the *Pgt* assembly (without phase switches) exhibited a clear phasing signal (Figure 4A). Over 145 Mb of the assembly have >80% of their Hi-C links to the same haplotype. The few *Pgt* contigs that have >80% of their Hi-C links to the opposite haplotype either belong to chromosomes 11A/11B that are unusual in that they both reside in the same nucleus in *Pgt* 21-0 (Li, Feng *et al.*, 2019), or are small contigs < 50 Kb. In contrast, the uncorrected *Pgt* assembly with phase switch contigs included has a smaller proportion of the assembly confidently assigned to the correct phase and noisy Hi-C links are clearly visible (Figure 4A).

**Figure 4:**
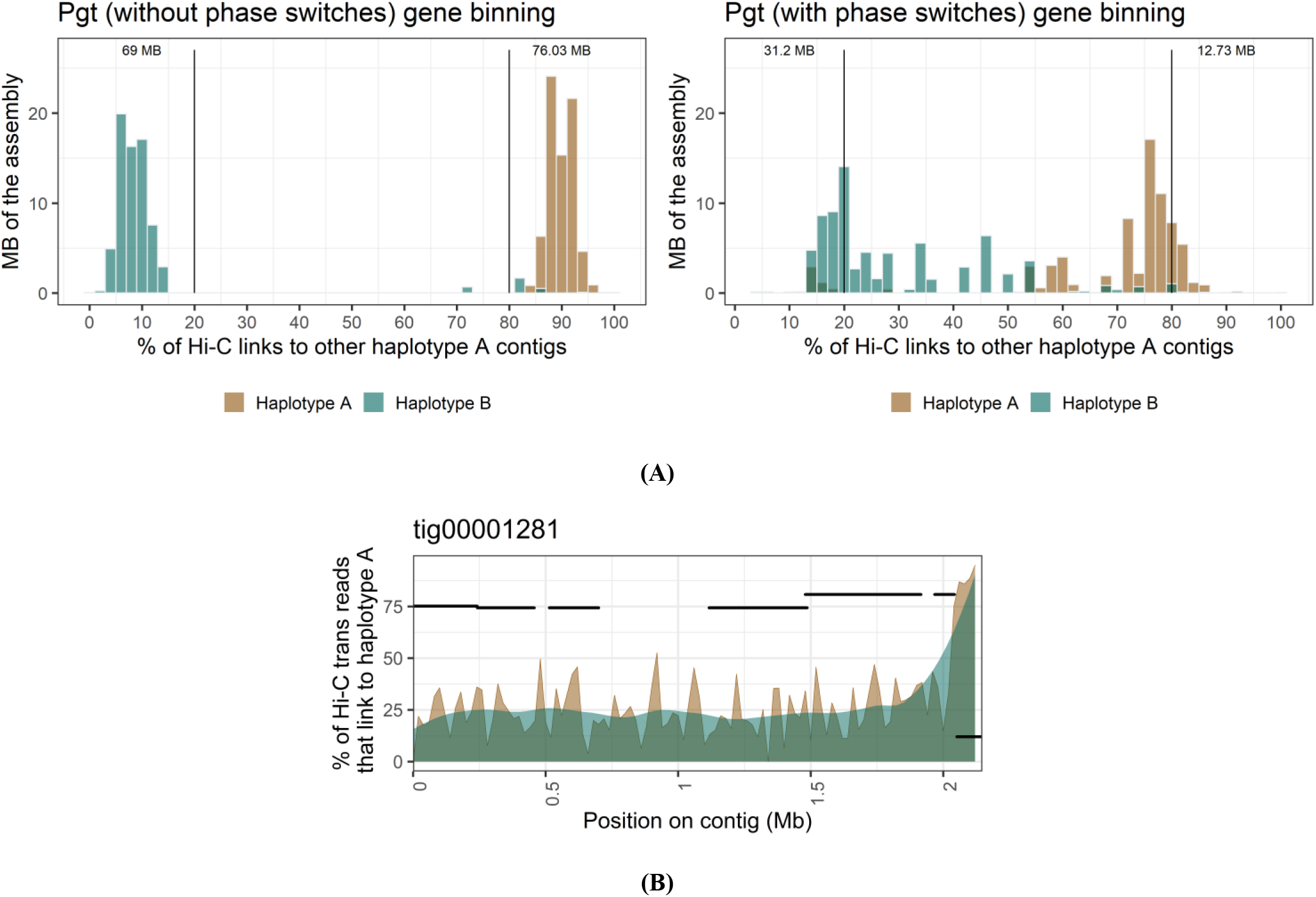
Hi-C links between the two haplotypes after the gene binning step for the *Pgt* assembly. **(A)** Hi-C links within scaffold bins are excluded. The phasing signal in the *Pgt* assembly is clearly visible when phase switches are corrected. Over ~69 Mb of each haplotype have more than 80% of their Hi-C contacts to the same haplotype. In contrast, the presence of phase switches causes a noisier phasing signal between the haplotypes. **(B)** *Pgt* **contig tig00001281 with its associated haplotigs (black segments).** The % of Hi-C trans contacts that link to haplotype A (with an associated smoothing line) are shown. Haplotigs are shown at the *y* coordinate that corresponds to their Hi-C contacts to haplotype A. This contig has a phase switch previously detected by parental data at ~2.062 Mb, which is also visible here with Hi-C data alone at the haplotig alignment boundary.

To address phase switch correction approaches, we visualized the proportion of Hi-C contacts to haploypes A and B for a contig that contains known phase switches. Previously, phase switches in *Pgt* were identified by using alignment to nucleus-specific contigs from a natural hybridisation event (Li, F. *et al.*, 2019). For example, *Pgt* contig tig00001281 was previously broken into two contigs at the genomic coordinate 2.062 Mb informed by haplotig alignments. This breakpoint also clearly stands out using Hi-C contact data (Figure 4B). Here, most Hi-C contacts switch from one haplotype to the other in that region and the corresponding haplotigs also switch phase at that point (Figure 4B). Thus, our Hi-C based method can be used to detect and correct phase switches at haplotig boundaries.

Next, we applied the phasing pipeline to the leaf rust assemblies. For the Nanopore assembly, the two haplotypes comprised 118.6 Mb and 83.8 Mb, respectively. The difference in size between these results from the absence of the collapsed sequence regions in one of the haplotypes. In contrast, the HiFi-Canu assembly returned the two haplotypes at 120.4 Mb and 120.1 Mb. The HiFi-hifiasm assembly phased into the two haplotypes with 119.3 Mb and 122.8 Mb. Whilst both the Nanopore and HiFi scaffold bins phased into exactly two communities essentially representing two haploid genome contents (with the collapsed regions only represented once in the Nanopore assembly), they exhibited major differences in Hi-C phasing signal strength. We found that the HiFi-Canu and HiFi-hifiasm exhibit similar phasing profiles to the *Pgt* raw assembly, again suggesting the presence of a few contigs with phase switches (Figure 5). In contrast, the Nanopore assembly did not show a clear phasing signal, with most contigs of haplotype A showing Hi-C links to haplotype B and vice versa. This suggested that the presence of extensive phase switches in this assembly precludes the accurate separation of haplotypes by this approach.

**Figure 5:**
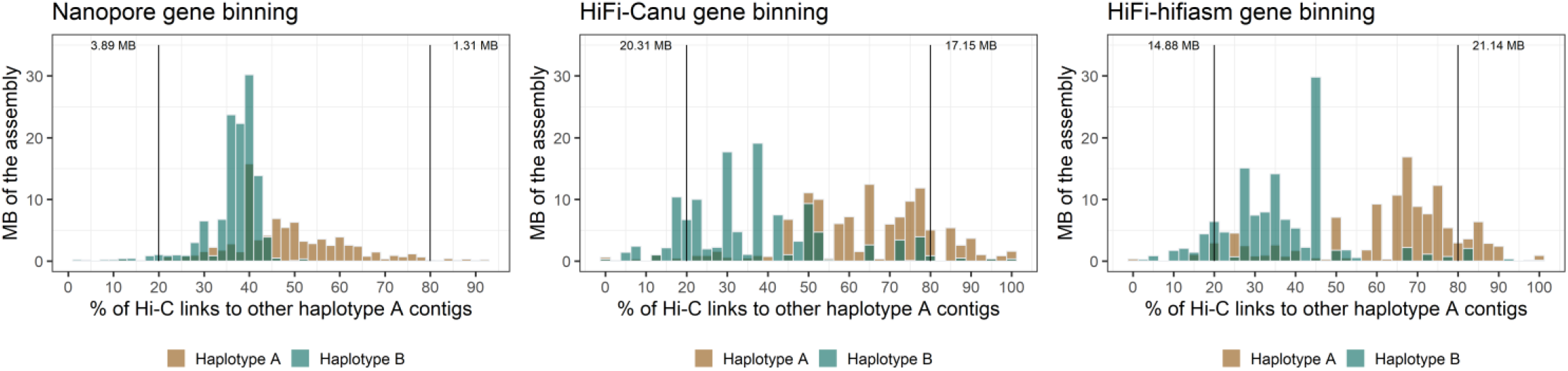
Hi-C links between the two haplotypes after the gene binning step for the *Pt76* assemblies. Hi-C links within scaffold bins are excluded. The phasing signal between scaffold bins in the Nanopore assembly is weak with only ~5 Mb of each haplotype have more than 80% of their Hi-C contacts to the same haplotype. This indicates the presence of extensive phase switches. In contrast, the HiFi assemblies show a phasing signal similarly to the *Pgt* assembly with phase switches present (Figure 4).

We investigated if phase switches cluster in large blocks of genomic regions or if they are randomly distributed along the contigs. For this, we visualized the proportion of Hi-C contacts to haploypes A and B for each scaffold bin. For example, scaffold bin *Bin_2_* in the HiFi-Canu assembly has two haplotype sets *Bin_2_a_* (8 contigs, total 4.43 Mb) and *Bin_2_b_* (contig tig00000828, 4.35 Mb). Contig tig00000828 appears to switch phase twice at ~1.5 and ~3.7 Mb, which coincides with the corresponding haplotig alignment start and end points (Figure 6A). This process identified phase switch sites in 17 contigs in the HiFi-Canu assembly, which were also supported by drops in Illumina read coverage. This allowed these contigs to be manually corrected by breaking at the switch site. The correction of these phase switches reduced the proportion of *trans*-contacts between haplotypes in the HiFi-Canu assembly from 8.3% to 3.8%, a similar rate to the benchmark fully-phased *Pgt* assembly (2.7%, Table 2). Similarly, we found and corrected 14 contigs with phase switches in the HiFi-hifiasm assembly, which decreased the proportion of *trans*-contacts between haplotypes from 7.3% to 3.4% (Table 2, Figure 6B). However, two large contigs (> 6 Mb) in the HiFi-hifiasm assembly appear to have phase switches that do not clearly align with the haplotig boundaries and we did not attempt to fix these potential errors (Supplementary Figure S2). Inspection of the graphs for the Nanopore assembly did not allow for identification of phase switch boundaries: firstly because they appeared to be more numerous; secondly they did not appear to correspond to haplotig boundaries; and thirdly the presence of collapsed regions with intermediate haplotype connection levels obscured the signal (Figure 6C). Therefore, we did not correct phase switches in the Nanopore assembly. In the following sections, we proceeded with the phase switch-corrected HiFi-Canu and the Nanopore assemblies for a comparison on the chromosome-scale level.

**Figure 6:**
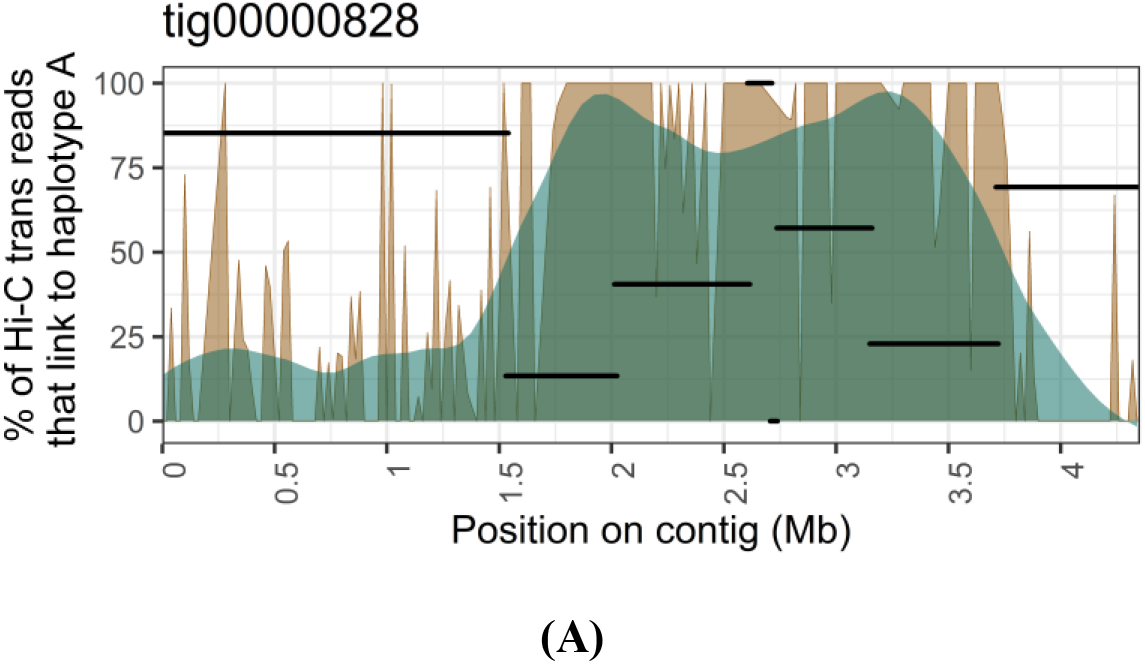
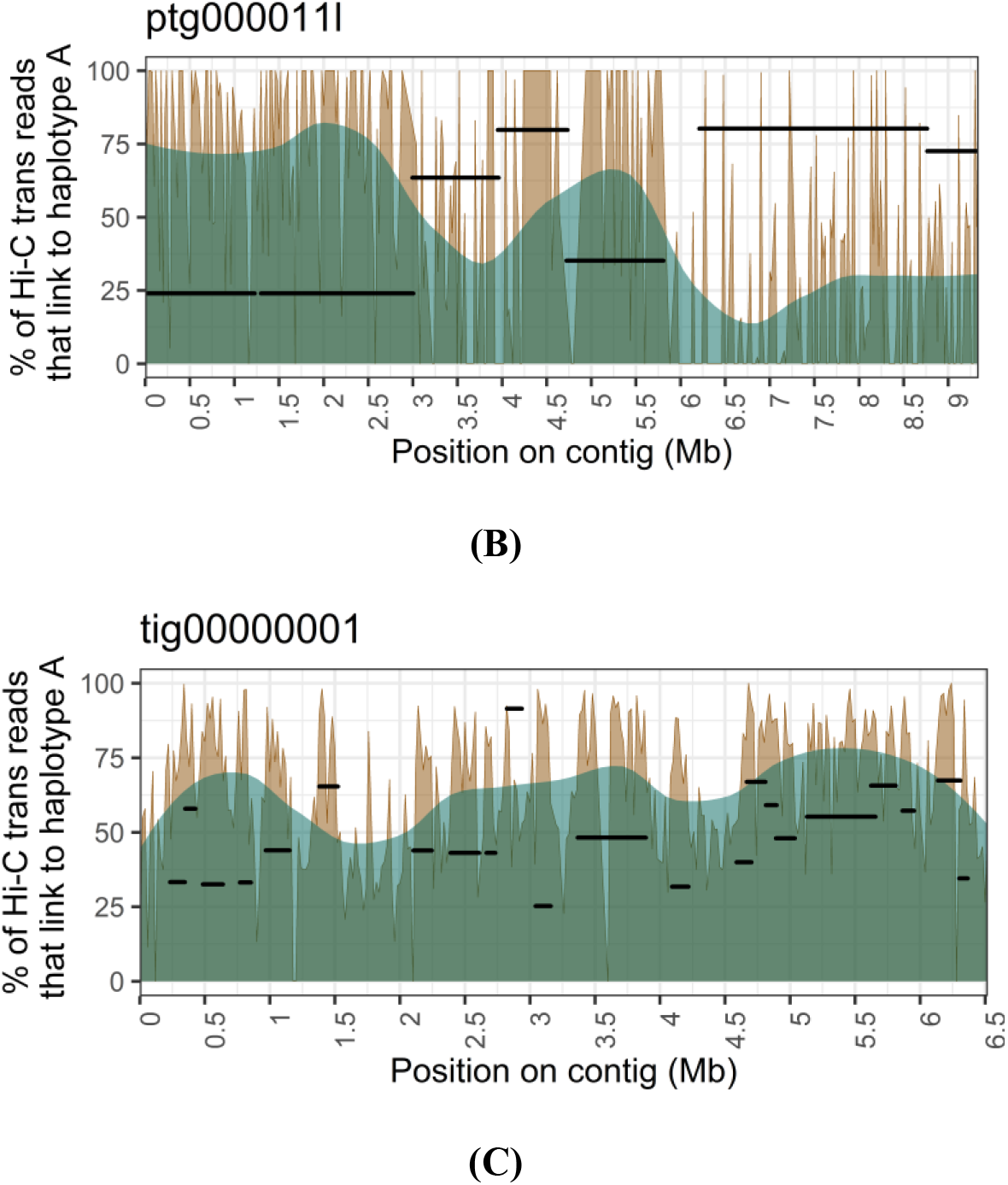
A contig with its associated haplotigs for each of the three *Pt76* assemblies (black segments). The % of Hi-C trans contacts that link to haplotype A (with an associated smoothing line) are shown. Haplotigs are shown at the *y* coordinate that corresponds to their Hi-C contacts to haplotype A. If a haplotig has no Hi-C contacts, it is shown at at *y* = 100. **(A)** Contig tig00000828 from the HiFi-Canu assembly and its associated haplotig alignments. Contig tig00000828 appears to switch phase at ~1.5-3.7 Mb, which overlaps with the corresponding haplotig alignment start and end points. **(B)** Contig ptg000011l from the HiFi-hifiasm assembly and its associated haplotig alignments. Contig ptg000011l appears to switch phase at ~3-4.7 Mb and at ~5.8 Mb, which overlaps with the corresponding haplotig alignment start and end points. **(C)** Contig tig00000001 from the Nanopore assembly and its associated haplotig alignments. Distinct phase switch blocks such as in the HiFi assemblies are not visible. Collapsed regions where the Hi-C contacts are in the 50% range are clearly visible in the Nanopore assembly.

### Haplotype phasing and chromosome curation of the HiFi-Canu and Nanopore assemblies

Prior to scaffolding of the two *Pt76* assemblies (Nanopore and phase switch-corrected HiFi-Canu), we conducted further contig phase assignment based on iterative application of the above process (Figure 2). For this, we assigned the unplaced contigs that were not part of the scaffold bins over multiple rounds based on synteny and Hi-C contacts to the two haplotypes (Figure 2). Again, we first validated this phasing pipeline on the phase switch corrected *Pgt* assembly. This resulted in two haplotypes at 90.0 Mb and 83.0 (3.9 Mb unphased). Only one small contig (33.3 kb) is in disagreement with the published phased genome of *Pgt* (Li, F. *et al.*, 2019). Our method also correctly captures the single chromosome exchange event in this isolate, where both chromosome 11 haplotypes are in the same nucleus. The final *Pgt* haplotypes have a strong phasing signal, with all contigs having more than 80% of their Hi-C contacts to other contigs in the same haplotype (Figure 7).

**Figure 7:**
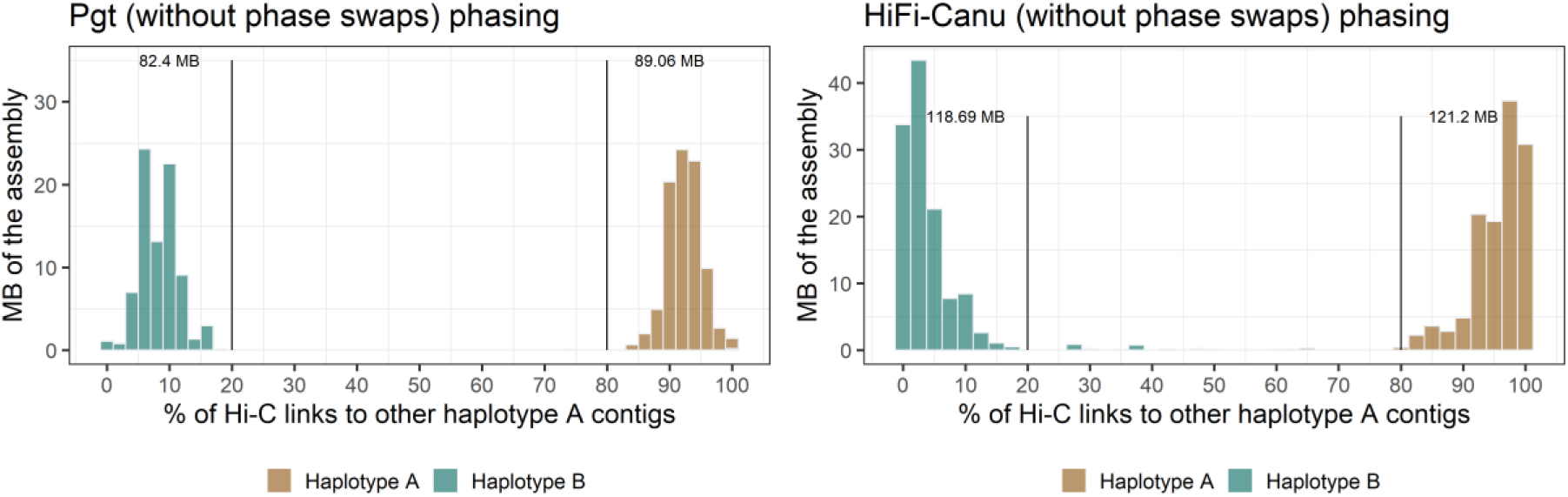
Hi-C links for the *Pt76* HiFi leaf rust assembly after the final phasing step. The phasing signal for the HiFi-Canu assembly is clearly visible and has a similar profile to the fully-phased *Pgt* assembly.

For the *Pt76* HiFi-Canu assembly, application of the same phasing pipeline yielded two haplotypes of 123.6 Mb and 122.1 Mb (10.7 Mb remained unphased). The correction of phase switches reduced the proportion of *trans*-contacts between haplotypes in the HiFi-Canu assembly from 8.3% to 3.8%, a similar rate to the gold-standard fully-phased *Pgt* assembly (3.2%). The phasing signal is evident in the HiFi-Canu assembly without phase switches (Figure 7). The Nanopore assembly could not be phased into the two haplotypes with our method. The extensive presence of phase switches leads to numerous Hi-C contacts between haplotypes. Whilst the Nanopore scaffold bins separated into two haplotypes after synteny assignment with a weak phasing signal (two haplotypes of 100.7 Mb and 120.7 Mb), subsequent assignment of unplaced contigs based on their Hi-C contacts over multiple rounds resulted in two haplotypes at 219.6 Mb and 1.8 Mb. This is due to the last quality control check in the pipeline, where the Hi-C contacts of all contigs are inspected and if a contig has over 50% of its Hi-C contacts to the other haplotypes, its assignment is swapped to appropriate haplotype. We did not run this final step for the Nanopore assembly before chromosome curation, only for the HiFi assemblies.

We curated pseudo-haplotype Nanopore chromosomes and fully-phased HiFi-Canu chromosomes by scaffolding the two haplotypes separately using Hi-C data and then further joined scaffolds into chromosomes through visual inspection of Hi-C contact maps. This resulted in 18 chromosomes for each haplotype, with 18 centromeres clearly visible in the Hi-C contact map of each haplotype (Supplementary Figure S3) as distinct outwards-spreading bowtie shapes previously described in *Pgt* (Sperschneider et al., 2020). The Nanopore chromosomes are 124.43 Mb and 106.95 Mb in length and the HiFi-Canu chromosomes are 123.9 Mb and 121.6 Mb in length (Table 3). Based on the estimated haploid genome size of ~132 Mb this suggests that some repetitive regions might not have been able to be scaffolded due to low Hi-C signal in those regions. Indeed, 11.0 Mb of mainly small contigs < 100 Kb (L50: 30.2 Kb) remained unplaced. However, the near-complete assembly of the gene space in the HiFi assembly is supported by the high BUSCO scores for each of the haplotypes at 95.5% and 95.2%, respectively. BUSCO scores are lower in the Nanopore assembly at 91 % for haplotype A and only 68% for haplotype B due to the absence of large collapsed regions in this haplotype. A long-read coverage analysis estimated that in the HiFi chromosomes only 1.6 Mb are collapsed genomic regions on haplotype A and 1.0 Mb are collapsed genomic regions on haplotype B. In contrast, 3.4 Mb are collapsed genomic regions in the unplaced contigs. We did not assign the unplaced contigs to a haplotype as the Hi-C signal is weak due to multimapping in these repetitive contigs.

**Table 3:** Assembly statistics for the *Pt76* chromosome assemblies. Assembly statistics for the Nanopore and HiFi haplotypes. Nanopore haplotype A has a high proportion of collapsed regions which are absent in haplotype B, leading to its smaller size. The HiFi haplotypes are of similar size and have a higher proportion of complete BUSCOs than the Nanopore haplotype A.

|  | Nanopore chromosomes |  |  | HiFi-Canu chromosomes |  |  |
| --- | --- | --- | --- | --- | --- | --- |
|  | Haplotype A | Haplotype B | Unplaced contigs | Haplotype A | Haplotype B | Unplaced contigs |
| Assembly size | 124.430 Mb | 106.954 Mb | 2.351 Mb | 123.9 Mb | 121.6 Mb | 11.0 Mb |
| # of scaffolds | 18 | 18 | 46 | 18 | 18 | 362 |
| Scaffold N/L50 | 8/7.596 Mb | 8/6.085 Mb | 12/81.914 Kb | 8/7.7 Mb | 8/7.4 Mb | 140/30.2 Kb |
| Max scaffold length | 9.699 Mb | 8.736 Mb | 139.988 Kb | 9.5 Mb | 9.3 Mb | 98.9 Kb |
| GC content | 46.64% | 46.61% | 44.43% | 46.6% | 46.7% | 42.4% |
| Complete BUSCOs (%) | 91.1% | 68.2% | 0.5% | 95.5% | 95.2% | 2.3% |
| Duplicated BUSCOs (%) | 2.6% | 2.2% | 0% | 3.7% | 4.2% | 0.1% |
| Fragmented BUSCOs (%) | 6.1% | 7.6% | 0.2% | 3.1% | 2.8% | 0.2% |
| Predicted genes | 14,601 | 12,549 | 256 | 14,482 | 13,552 | 571 |

Over 99% of the Nanopore chromosome assembly is represented in the HiFi chromosome assembly (Table 4). However, the HiFi chromosome assembly contains 2.5% of sequence that is not represented in the Nanopore chromosome assembly. The *Pt76* haplotypes are more similar than the *Pgt* haplotypes (Table 4). In *Pgt,* the average identity of aligned bases is ~95% (*Pt76*: ~99%) and large structural variation occurs (~12% unaligned bases; *Pt76*: ~2% unaligned bases). Annotation of both assemblies yielded similar gene content. The HiFi chromosomes have 14,482 and 13,552 predicted genes on haplotype A and B, respectively, compared to 14,601 predicted genes on Nanopore haplotype A. This is a reduction in gene content compared to the close relative *Pgt*, which has ~18,500 genes on each haplotype and a haploid genome size of only ~88 Mb (Li, F. *et al.*, 2019). Repeat prediction shows the expansion in the *Pt* genome size is almost entirely due to increased repetitive sequence content (151 Mb compared to 74 Mb), particularly retroelements and unclassified repeats (Table 5).

**Table 4:** Comparisons between the Nanopore and HiFi chromosomes as well as between the haplotypes of *Pt76* and *Pgt*.

|  | Comparison between *Pt76* chromosomes |  | Comparison between HiFi-Canu *Pt76* chromosomes |  | Comparison between *Pgt* chromosome assemblies |  |
| --- | --- | --- | --- | --- | --- | --- |
|  | HiFi-Canu | Nanopore | Haplotype A | Haplotype B | Haplotype A | Haplotype B |
| Number of sequences | 36 | 36 | 18 | 18 | 18 | 18 |
| Aligned bases | 97.6% | 99.1% | 97.1% | 98.9% | 85.6% | 89.5% |
| Unaligned bases | 2.5% | 0.9% | 2.9% | 1.1% | 14.4% | 10.5% |
| Average identity of 1-to-1 alignments | 99.7% |  | 99.5% |  | 95.8% |  |
| Average identity of M-to-M alignments | 99.7% |  | 99.0% |  | 95.4% |  |
| Translocations | 963 | 1,023 | 1,161 | 1,179 | 12,087 | 12,139 |
| Inversions | 51 | 49 | 163 | 174 | 537 | 566 |
| Insertions | 1,732 | 3,206 | 10,896 | 11,548 | 40,159 | 49,047 |
| Tandem duplication insertion | 12 | 64 | 329 | 346 | 86 | 121 |
| Total SNPs | 62,461 |  | 334,571 |  | 1,420,848 |  |
| Total Indels | 249,485 |  | 186,609 |  | 877,171 |  |

**Table 5:** Predicted repeat content of the *Pt76* and *Pgt* chromosomes. The *Pt76* chromosomes are larger in size and this is driven by repeat expansion, particularly of retroelements and unclassified repeats.

|  | *Pt* HiFi chromosomes | *Pgt* chromosomes |
| --- | --- | --- |
| Number of scaffolds | 36 | 36 |
| Total length | 244.8 Mb | 169.9 Mb |
| GC content | 46.6% | 43.5% |
| Bases masked | 62.1% | 44.4% |
| Retroelements (% of sequence) | 18.9% | 12.9% |
| Ty1/Copia (% of sequence) | 5.9% | 3.1% |
| Gypsy/DIRS1 (% of sequence) | 11.0% | 8.6% |
| DNA transposons (% of sequence) | 6.4% | 5.5% |
| Unclassified (% of sequence) | 35.2% | 24.5% |

### HiFi sequencing can be used to phase genomes with ~0.7% heterozygosity with ~30-40x haploid genome coverage

Comparison of the two haplotypes in the *Pt76* assembly showed that at least 97.3% of bases could be aligned with an average identity of ~99% (Table 4). This is supported by an Illumina *k*-mer analysis using GenomeScope which indicates a heterozygosity rate of 0.7% (Vurture *et al.*, 2017). This is a considerably lower divergence between haplotypes than observed for *Pgt*, where as low as 85% of the haplotypes could be aligned with identities of only ~95% in aligned regions. This indicates that the Hifi sequence data was able to reliably separate the haplotypes even with relatively low inter-haplotype divergence. To investigate the effect of HiFi read coverage on assembly contiguity and haplotype separation, we downsampled the HiFi reads to various levels of haploid genome coverage and assembled them again using Canu (Table 6). As expected, the assemblies improved in contiguity with increased coverage (Table 6). At least 25x coverage seems to be required for achieving L50 over 1 Mb, and the optimum assembly contiguity was at 40x coverage. We used Illumina mappings to the phased HiFi chromosomes to assess the rate of phase switches in the downsampled assemblies. For this, we mapped the Illumina reads to both of the HiFi chromosome haplotypes and then classified each read based on which haplotype it aligned to best (Total mapped reads: 63 million; haplotype A-specific: 5.2 million reads; haplotype B-specific: 4.9 million reads). Then we aligned these haplotype-specific reads to the each of the down-sampled assembly contig sets and recorded the proportion of the contigs covered by reads derived from each haplotype. In contigs without phase switches, we expect to see high coverage from reads specific to one haplotype and very low coverage from reads specific to the other haplotype. Indeed the phase-corrected HiFi-Canu assembly showed a very strong signal for correctly phased contigs, while the uncorrected version showed a small number of phase switch contigs with intermediate haplotype signals (Figure 8). HiFi assemblies below 30x genome coverage have a large proportion of contigs with phase switches and genome coverage of 40x appears to be the assembly with the strongest phasing signal, and equivalent to the original 50x assembly (Figure 8). This suggests that low coverage HiFi data is not suitable for achieving fully-phased assemblies and that there is also a high coverage limit beyond which phasing of the assembly does not improve.

**Table 6:** Statistics for HiFi assemblies at various levels of haploid genome coverage. Genome coverage of at least 25x is required to achieve a contiguous assembly. However, assemblies below 30x coverage have a large proportion of contigs with phase switches (Figure 8).

| Statistic | HiFi-Canu assemblies |  |  |  |  |  |
| --- | --- | --- | --- | --- | --- | --- |
| Haploid genome coverage | 50x | 40x | 30x | 25x | 20x | 10x |
| Assembly size | 269.0 Mb | 267.5 Mb | 258.1 Mb | 254.5 Mb | 252.3 Mb | 212.5 Mb |
| # of contigs | 1,063 | 982 | 825 | 872 | 1,215 | 3,285 |
| Assembly N/L50 | 28/2.4 Mb | 28/2.7 Mb | 48/1.7 Mb | 72/1.1 Mb | 139/493.9 Kb | 735/86.4 Kb |
| Assembly N/L90 | 176/119.6 Kb | 162/141.0 Kb | 220/186.0 Kb | 307/153.4 Kb | 566/93.6 Kb | 2379/30.5 Kb |
| Maximum contig length | 9.3 Mb | 10.6 Mb | 5.8 Mb | 5.1 Mb | 2.7 Mb | 563.8 Kb |

**Figure 8:**
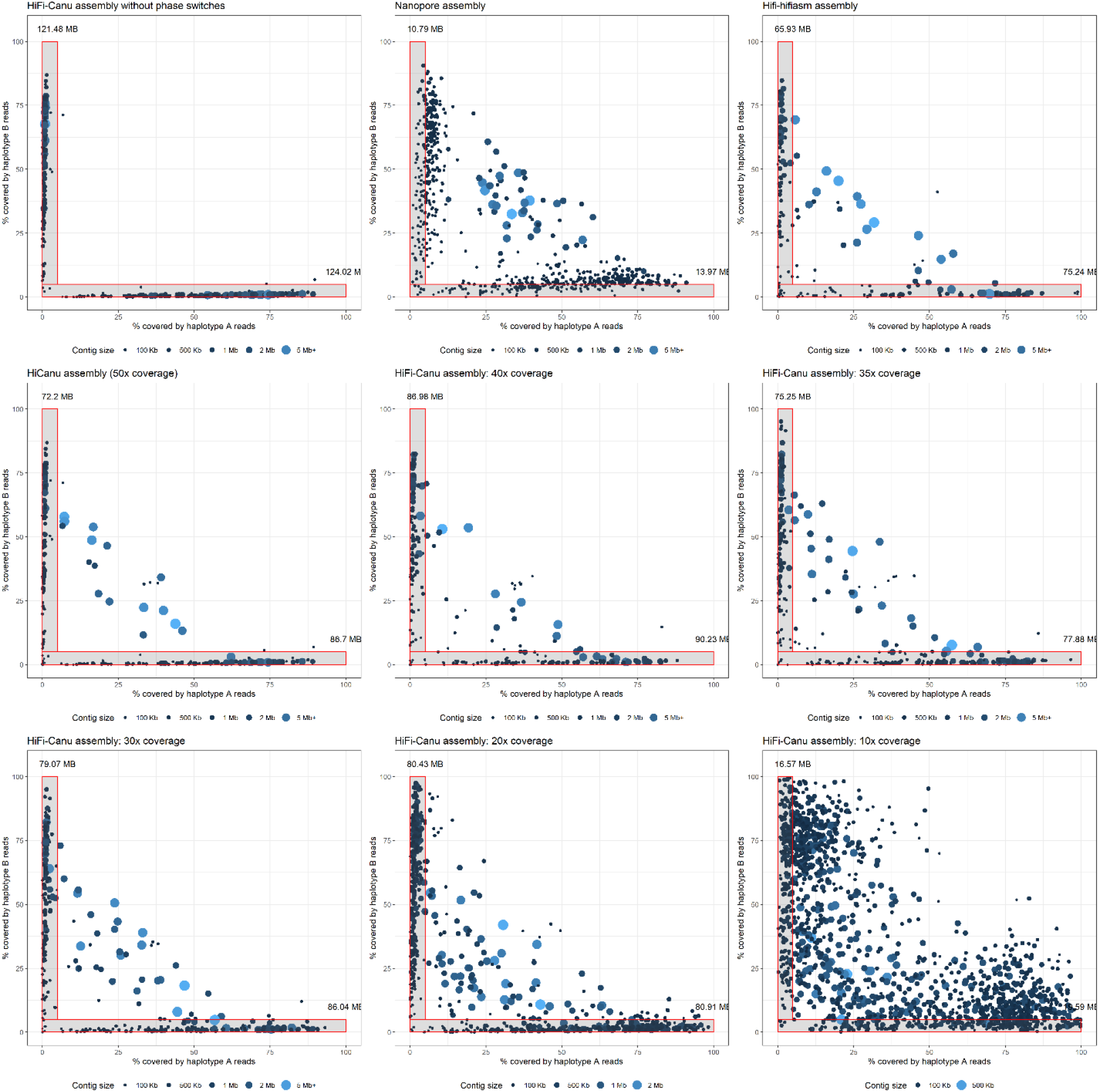
Contig coverage of haplotype-specific reads for HiFi assemblies at varying levels of genome coverage. Illumina reads were binned into haplotype-specific reads if they align better to one of the haplotypes from the HiFi chromosomes. These haplotype-specific reads were then re-mapped to the contigs of the downsampled HiFi assemblies as well as to the Nanopore and HiFi-hifiasm assembly. The proportion of each contig covered by reads specific to haplotypes A and B is shown. The HiFi assembly at 40x genome coverage has the clearest phasing signal and assemblies below 30x coverage have many phase switches.

## Discussion

Long-read sequencing together with scaffolding data (Hi-C, optical maps or genetic linkage maps) is the foundation for achieving chromosome-scale assemblies across a wide range of species. However, phasing of non-haploid genomes is still a challenging problem. Even with highly accurate or ultra-long reads such as HiFi or Nanopore, genome assemblers will output some incorrectly assembled contigs with phase switches or chimeric misjoins. Scaffolding programs have limited ability to detect and correct these contigs and when applied to an unphased assembly will return incorrectly joined scaffolds that are an artificial mix of the haplotypes. To overcome this, several strategies have been proposed in diploid genomes such as human. First, when parental sequencing data is available, trio binning can first partition long reads from an offspring into haplotype-specific sets for subsequent assembly (Koren *et al.*, 2018). However, parental data is not always obtainable and genomic regions that are heterozygous in the parents cannot be phased. Second, some assemblers use *k*-mers or short reads from Hi-C or strand sequencing data to partition HiFi reads into the haplotypes (Edge *et al.*, 2017; Garg *et al.*, 2021; Porubsky *et al.*, 2021). These methods can struggle to identify haplotype-specific variants in complex or collapsed genomic regions (Cheng *et al.*, 2021). Falcon-Phase is a recent Hi-C based method for phasing that re-assigns genomic regions that have haplotigs to its phase (Kronenberg *et al.*, 2021). Falcon-Phase is currently based on the output from a Falcon-Unzip assembly and will automatically backfill homozygous regions in its pseudo-haplotype output. Methods that are applicable to genome assemblies of polyploid plant species or fungal species that carry multiple haploid nuclei per cell were thus far still lacking.

We showed that genome assemblies both from Nanopore or HiFi sequencing reads with either the Canu or hifiasm assemblers will contain contigs with chimeric misjoins or phase switches. To achieve a high-quality chromosome-scale haplotype-phased assembly, these errors must be detected and corrected. When Hi-C data is available, chimeric misjoins between different chromosomes are clearly visible in Hi-C contact maps and can be manually corrected. In contrast, we found that phase switches are not clearly visible in Hi-C contact maps for the assembly. However, we showed that through first phasing a reliable subset of the haplotypes through gene binning and then visualizing the Hi-C contacts to each haplotype across the contigs, these phase switch errors can be detected. We envision that our Hi-C contact graph method can be extended to polyploid plant genomes in the future. Whilst haplotypes in polyploid plants do not reside in separate nuclei, the Hi-C contact signal within a chromosome will be stronger than between chromosomes and this could be used in our phase switch detection approach. This is a similar concept to ALLHiC, which builds a Hi-C contact graph and ignores Hi-C signal between allelic contigs (Zhang *et al.*, 2019). However, for ALLHiC the number of chromosomes needs to be provided by the user and phase switches cannot be corrected during the assembly process. In contrast, our method provides a framework for phase switch identification and correction, which is the foundation for fully-phased genomes.

We showed how Hi-C data alone can be used in dikaryons to fully phase the haplotypes and achieve nuclear-resolved chromosome-scale assemblies. However, the input genome assembly quality is crucial. Highly collapsed assemblies or assemblies with a very high number of phase switches will not be able to be phased. HiFi data has been reported to be able to separate haplotypes up to a divergence of 0.01% with appropriate genome coverage. Whilst the highly heterozygous *Pgt* was able to be assembled into the two haplotypes with PacBio RSII long read sequencing (Li, F. *et al.*, 2019), HiFi sequencing data is likely essential to achieve haplotype separation in the *Pt76* assembly that is ~0.7% heterozygous. Furthermore, we showed that for *Pt76*, HiFi genome coverage of ~30-40x is required for producing an assembly that can be confidently phased. This is in line with current practice by the Canu assembler which downsamples HiFi read sets to 50x coverage by default.

We deliver the second fully-phased assembly for a rust fungus and a substantial improvement over previous leaf rust assemblies. The previously published *Pt104* assembly (Wu *et al.*, 2020) has an assembly size of 140.5 Mb, however it also contains 12.2% duplicated BUSCOs and is thus an overestimation of the true haploid genome size. In that case, PacBio Sequel data was used to generate a Falcon-Unzip assembly (Chin *et al.*, 2016). The primary assembly has L50 of 2.1 Mb (162 contigs, 92% complete BUSCOs), compared to the haplotig set with L50 of 816 Kb (713 contigs, 84% complete BUSCOs). Whilst this fragmentation of the haplotigs could be an artefact of the assembler used, it might also be due to the error rate of the Sequel technology that is higher than for HiFi sequencing reads.

The critical feature required for phase separation between haplotypes is sequence accuracy. We showed that relatively low-coverage MinION Nanopore sequencing (~50x haploid genome coverage) is not appropriate for achieving haplotype-phased assemblies in the case of leaf rust. Obtaining higher coverage will likely improve the phase switch error rate in the Nanopore assembly through higher accuracy in the read correction step of the assembler. However, major advances in Nanopore sequencing will come from improved technology such as new pore types and sequencing chemistries as well as higher accuracy of base calling algorithms. In the future, we expect Nanopore sequencing accuracy to improve, in which case the longer read lengths may offer an advantage for fully resolving complex repetitive regions and assembly contiguity.

## Conclusions

Chromosome-scale haplotype reconstruction is essential for understanding genome evolution, pathogenicity and is the foundation for downstream comparative analysis. Here, we deliver the first Hi-C based phasing pipeline for dikaryons and compare HiFi to Nanopore technologies for accurate genome assembly. We highlight the importance of identifying phase switches in contigs and show that, in the absence of parental data, this can be achieved with Hi-C data alone. Our work highlights that current low-coverage Nanopore sequencing technology delivers a pseudo-haplotype representation of the genome, whereas HiFi sequencing delivers an assembly with relatively few phase switches. Further technological advances in Nanopore and PacBio sequencing will lay the foundation for a new era of gapless end-to-end, fully-phased assemblies in species that have previously been overlooked. This will lay the foundation for understanding of genome evolution and other biological phenomena.

## Materials and Methods

### Sampling and pathotyping of *Puccinia triticina* isolate *Pt76*

Rust infected samples from the leaf rust susceptible wheat cultivar *Morocco* were collected from the CSIRO field site in Canberra during November 2018. A *Puccinia triticina* (*Pt*) culture was purified through single pustule isolation and pathotyped using the standard wheat differential sets carrying unique resistance genes for leaf rust as described in the Cereal Rust Report, PBI. Based on the phenotypic resistance response, the isolate belongs to the 76-3,5,7,9,10,12,13 pathotype and was named *Pt76*.

### Oxford Nanopore Technologies native long-read DNA sequencing

Urediniospores were snap frozen in liquid nitrogen and stored at −80°C until DNA extraction. High-molecular weight DNA was extracted from the spores using a modified Cetrimonium bromide (CTAB) extraction protocol which described in detail on Protocols.io (dx.doi.org/10.17504/protocols.io.5isg4ee). Briefly, approximately 600 mg of spores were homogenised to a fine powder using a mortar and pestle, which was kept frozen with liquid nitrogen. The sample was then incubated with a CTAB based lysis buffer with RNAse A and Proteinase K. The mixture was then cleaned with chloroform:isoamyl alcohol (24:1, v/v), transferring the upper phase to a CTAB precipitation buffer. After 1 h, white crystals of CTAB-DNA complexes formed, which were pelleted by centrifugation. The supernatant was discarded, and the pellet washed with 70% ethanol multiple times. After air drying, the pellet was resuspended in nuclease-free water and stored at 4°C. This crude DNA underwent size selection for fragments ≥ 20 kb using a PippinHT (Sage Science) prior to sequencing. An Oxford Nanopore Technologies long-read native DNA sequencing library was prepared according to the manufacturer’s protocol 1D genomic DNA by ligation (SQK-LSK109). Sequencing was performed on a MinION Mk1B using a FLO-MIN106 R9.4.1 revD flow cell, according to the manufacturer’s instructions. MinION fast5 reads were basecalled to fastq with Guppy version 4.0.11. Sequencing output and quality was inspected with NanoPlot version 1.28.2 (De Coster *et al.*, 2018).

### PacBio HiFi DNA sequencing

High molecular DNA from urediniospores was extracted as previously described (Li, F. *et al.*, 2019). DNA quality assessed with a Nanodrop Spectrophotometer (Thermo Scientific, Wilmington, DE, USA) and the concentration quantified using a broad-range assay in Qubit 3.0 Fluorometer (Invitrogen, Carlsbad, CA, USA). DNA library preparation (10-15 Kb fragments Pippin Prep) and sequencing in PacBio Sequel II Platform (One SMRT Cell 8M) were performed by the Australian Genome Research Facility (AGRF) (St. Lucia, Queensland, Australia) following manufacturer’s guidelines.

### Illumina short-read whole-genome and Hi-C DNA sequencing

Multiplexed, short-read, whole-genome DNA sequencing libraries were generated using a cost-optimized, transposase-based protocol (dx.doi.org/10.17504/protocols.io.unbevan), based on Illumina Nextera XT DNA Library Prep (Document # 15031942 v03 February 2018). Chromosome conformation was captured and a sequencing library prepared using a Microbe Proximo Hi-C Kit from Phase Genomics, according to the manufacturer’s ProxiMeta™ Hi-C Protocol (version 1.5, 2019). However, further action was taken to ensure fungal cell lysis, by adding a 3mm ball bearing to the tube grinding with a TissueLyser II (Qiagen). Sequencing libraries underwent size selection for fragments with insert sizes of 300-500 bp using a PippinHT (Sage Science). Illumina short-read sequencing was performed on a NextSeq 500 using a mid-output 300 cycles flow cell (150 bp paired-end, 130 million clusters).

### Genome assembly and polishing

For the Nanopore assembly, Canu 2.0 (Koren *et al.*, 2017) was run with the parameters genomeSize=120m corOutCoverage=200 “batOptions=-dg 3 -db 3 -dr 1 -ca 500 -cp 50”. The assembly was polished once with Racon 1.4.13 (-m 8 -x −6 -g −8 -w 500 --no-trimming --include-unpolished) (Vaser *et al.*, 2017) once with medaka 1.0.3 (-v and model r941_min_high_g360, https://nanoporetech.github.io/medaka/) and twice with Pilon 1.22 (Walker *et al.*, 2014) (--fix indels). For the HiFi assemblies, Canu 2.0 (Koren *et al.*, 2017) was run with the parameters genomeSize=120m and -pacbio-hifi and hifiasm 0.13 was run with default parameters (Cheng *et al.*, 2021).

### Cleaning of the assemblies

Hi-C contact maps were produced with HiC-Pro 2.11.1 (Servant *et al.*, 2015) (MAPQ=10) and visually examined for the presence of mis-assemblies and chimeric contigs. Breakpoints for chimeric contigs were identified through visual inspection of contact maps and long-read alignments to the contigs with minimap2 (Li, 2018) and the flag -- secondary=no. Contaminants were identified using sequence similarity searches (BLAST 2.9.0 -db nt -evalue 1e-5 - perc_identity 75) (Altschul *et al.*, 1990) in combination with sequence coverage and GC content analysis. For sequence coverage, we aligned the long reads to the polished assembly with minimap2 (Li, 2018) and the flag --secondary=no. GC content and coverage was called using bbmap’s pileup.sh tool on the minimap2 alignment file (http://sourceforge.net/projects/bbmap/). Combining the sequence similarity search results and GC content, some contigs were identified to be of bacterial origin. The mitochondrial contig was identified based on BLAST searches against a mitochondrial database. Several small contigs were identified as high-coverage duplicated fragments of the primary mitochondrial contig. All contaminant contigs and the mitochondrial contigs were removed from the assembly. Collapsed regions were determined with a long-read mapping and coverage analysis (https://github.com/JanaSperschneider/GenomeAssemblyTools/tree/master/CollapsedGenomicRegions).

### SNP calling

Illumina reads were aligned to the assemblies with BWA-MEM 0.7.17 (Li & Durbin, 2009) and duplicate reads were marked with sambamba markdup (Tarasov *et al.*, 2015). FreeBayes 1.3.2 was run with --ploidy 2 (Garrison & Marth, 2012) and vcftools 0.1.16 was used to filter SNPs with options --minQ 30 –recode (Danecek *et al.*, 2011).

### Gene binning and phasing method

For the clean assemblies, a table of BUSCO gene hits (BUSCO 3.1.0 -l basidiomycota_odb9 -m geno -sp coprinus) (Simao *et al.*, 2015) was produced as well as a table of gene hits from the *Puccinia triticina* BBBD race 1 transcript set (Cuomo *et al.*, 2017) with biokanga blitz (4.4.2 --sensitivity=2 --mismatchscore=1) (https://github.com/csiro-crop-informatics/biokanga). Only genes that have exactly two hits to the assembly were retained as phasing markers. All-versus-all contig alignments were computed with minimap2 (-k19 -w19 -m200 -DP -r1000) (Li, 2018). We used the duplicated gene information to put contigs that share genes into scaffold bins. For each possible pair of contigs, we recorded the number of their shared genes (number of shared BUSCO genes + number of shared leaf rust genes). The total number of shared genes was normalized to shared gene density per Mb. We then constructed a graph where each contig is a node and a pair of contigs is connected by a weighted edge comprising the shared gene density. Two contigs were connected by an edge if their shared gene density per Mb is greater than 30, if they share more than two genes and if one of the contigs has more than 20% of its bases align with the other. A graph network approach was then used to find connected communities (Python’s NetworkX community.best_partition) (Blondel *et al.*, 2008). These connected communities represent scaffold bins that contain homologous pairs of sequences from each haplotype. Scaffold bins that contain contigs with a combined size > 1 Mb were kept and for each scaffold bin, the contigs within were separated into the two haplotype sets.

We produced a Hi-C contact map in ginteractions format with HiC-Pro 2.11.1 (MAPQ=30) (Servant *et al.*, 2015) and hicexplorer 3.6 (Ramirez *et al.*, 2018). The scaffold bins were then phased into the two haplotypes using Hi-C links between scaffold bins, but not within scaffold bins as these are likely spurious contacts between homologous sequences that reside in separate nuclei. For two scaffold bins *x* and *y*, the number of normalized Hi-C contacts between the bins were recorded from the contact map at 20,000 bp resolution. A graph was generated using the nodes *x_a_*, *x_b,_ y_a_* and *y_b_*. If the two scaffold bins have the same haplotype configuration, *x_a_* to *y_a_* and *x_b_* to *y_b_* should have the highest Hi-C contact frequency. Alternatively, if the two scaffold bins have opposite haplotype configuration, *x_a_* to *y_b_* and *x_b_* to *y_a_* should have the highest Hi-C contact frequency. We generated a graph between the haplotype sets with the following weighted edges: *x_a_* to *y_a_* are assigned the weight (Hi-C contact frequencies between *x_a_* to *y_a_* and *x_b_* to *y_b_*)/(Hi-C contact frequencies between *x* and *y*); *x_b_* to *y_b_* are assigned the weight (Hi-C contact frequencies between *x_a_* to *y_a_* and *x_b_* to *y_b_*)/(Hi-C contact frequencies between *x* and *y*); *x_a_* to *y_b_* are assigned the weight (Hi-C contact frequencies between *x_a_* to *y_b_* and *x_b_* to *y_a_*)/(Hi-C contact frequencies between *x* and *y*); *x_b_* to *y_a_* are assigned the weight (Hi-C contact frequencies between *x_a_* to *yb* and *x_b_* to *ya*)/(Hi-C contact frequencies between *x* and *y*). A graph network approach was then used to find connected communities (Python’s NetworkX community.best_partition) (Blondel *et al.*, 2008) and this returned two communities that represent the two haplotypes. The remaining unphased contigs that were not part of the scaffold bins were then assigned based on synteny with sequence alignments (minimap2 -k19 -w19 -m200 -DP -r1000). If a contig shares synteny (> 75% aligned bases) with a contig from one of the haplotypes, it was put into the opposite haplotype bin. Two rounds of synteny assignment were run to place contigs into haplotypes. Then, the remaining unphased contigs were assigned to haplotype bins based on Hi-C contact frequencies. If a contig shares has more than 20 Hi-C contacts to the haplotypes and if over 80% of these Hi-C contacts are with one of the haplotypes, they are assigned to be part of that haplotype. This process was run two times and followed by two more rounds of synteny assignment to place the remaining contigs. As a last quality control check, the Hi-C contacts of all contigs were inspected and if a contig has over 50% of its Hi-C contacts with the other haplotypes, its assignment was swapped to appropriate haplotype. The gene binning and phasing method is available at https://github.com/JanaSperschneider/NuclearPhaser.

### Chromosome scaffolding and comparisons

For scaffolding, the Hi-C reads were first mapped to each haplotype using BWA-MEM 0.7.17 (Li & Durbin, 2009). Alignments were then processed with the Arima Genomics pipeline (https://github.com/ArimaGenomics/mapping_pipeline/blob/master/01_mapping_arima.sh). Scaffolding was performed using SALSA 2.2 (Ghurye *et al.*, 2017). Chromosomes were compared to each other with mummer 4.0.0b2, using nucmer and dnadiff (Marcais *et al.*, 2018).

### RNA sequencing, gene prediction and repeat annotation

Total RNA from dormant urediniospores, germinated urediniospores and from rust infected leaves at 6 and 9 days post inoculation (dpi) was extracted using the Promega Maxwell® RSC Plant RNA Kit with a Maxwell® RSC instrument (Promega.com.au). The spores were induced for germination by placing on the surface of milli-Q water, at 100% humidity and 22°C conditions for 16 hours. Three biological replicates were maintained for each sample. NanoDrop™ spectrophotometer immediately following extraction. Approximately 20 μg of RNA in nuclease-free water was transferred to RNAstable tubes, supplied by GENEWIZ (www.genewiz.com), incubated at room temperature for five minutes, then mixed by pipetting. Samples were dried completely by SpeedVac for 1.5 h, then sent to the GENEWIZ Genomics Centre in Suzhou, China for RNA sequencing (RNAseq) using Illumina NovaSeq platform with 150 bp paired end configuration.

RNAseq reads were cleaned with fastp 0.19.6 using default parameters (Chen *et al.*, 2018). The repeatmasked genome was used for gene annotation. RNAseq reads were aligned to the genome with HISAT2 (version 2.1.0 --max-intronlen 3000 -- dta) (Kim *et al.*, 2019). Genome-guided Trinity (version 2.8.4 --jaccard_clip --genome_guided_bam -- genome_guided_max_intron 3000) was used to assemble transcripts (Grabherr *et al.*, 2011). Funannotate (version 1.7.4) was then used for gene prediction. First, funannotate train was run with the Trinity transcripts. De novo repeats were predicted with RepeatModeler 2.0.0 and the option -LTRStruct (Flynn *et al.*, 2020). These were merged with the RepeatMasker repeat library and RepeatMasker 4.1.0 was run with this combined repeat database (http://www.repeatmasker.org). Second, funannotate predict was run on the repeat-masked genome with options --ploidy 2 --optimize_augustus and weights: codingquarry:0. Previously published leaf rust ESTs were provided to funannotate with --transcript_evidence (Xu *et al.*, 2011). Third, funannotate update was run (--jaccard_clip).

### Low-coverage HiFi genome assemblies and phasing assessment

We downsampled the HiFi sequencing reads with seqtk sample (version 1.2, https://github.com/lh3/seqtk). Canu 2.0 (Koren *et al.*, 2017) was run on the downsampled sets with the parameters genomeSize=120m and -pacbio-hifi. We aligned the Illumina sequencing reads to the HiFi chromosome haplotypes separately with BWA-MEM 0.7.17 (Li & Durbin, 2009). We then classified them based on which haplotype they align to best using the alignment score (AS). We aligned the haplotype-specific reads to the assemblies with BWA-MEM 0.7.17 and kept only alignments with edit distance of zero. We then recorded contig coverage of haplotype-specific read alignments with bbmap’s pileup.sh (version 38.37 https://sourceforge.net/projects/bbmap/).

## Declarations

### Ethics approval and consent to participate

Not applicable.

### Consent for publication

Not applicable.

### Availability of data and materials

Sequencing reads are deposited under the NCBI Bioproject PRJNA725323 (https://www.ncbi.nlm.nih.gov/bioproject/725323) and additionally HiFi sequencing reads are deposited in the CSIRO Data Access Portal under the persistent link https://doi.org/10.25919/xbqb-px51. The phasing pipeline is available under https://github.com/JanaSperschneider/NuclearPhaser.

### Authors’ contributions

JS, BS, SP, MF and PND conceived the study and designed experiments. JS analysed and interpreted all data sets and wrote the manuscript. HD and TH performed genome assemblies and analysed data sets. AWJ, AS, RM, NMU and DL performed sequencing experiments. SP, AM and YH performed pathotyping and RNA sequencing experiments. All authors contributed to data analysis and manuscript writing. All authors read and approved the final manuscript.

## Acknowledgements

Not applicable.

## Supplementary Figures

**Supplementary Figure S1:**
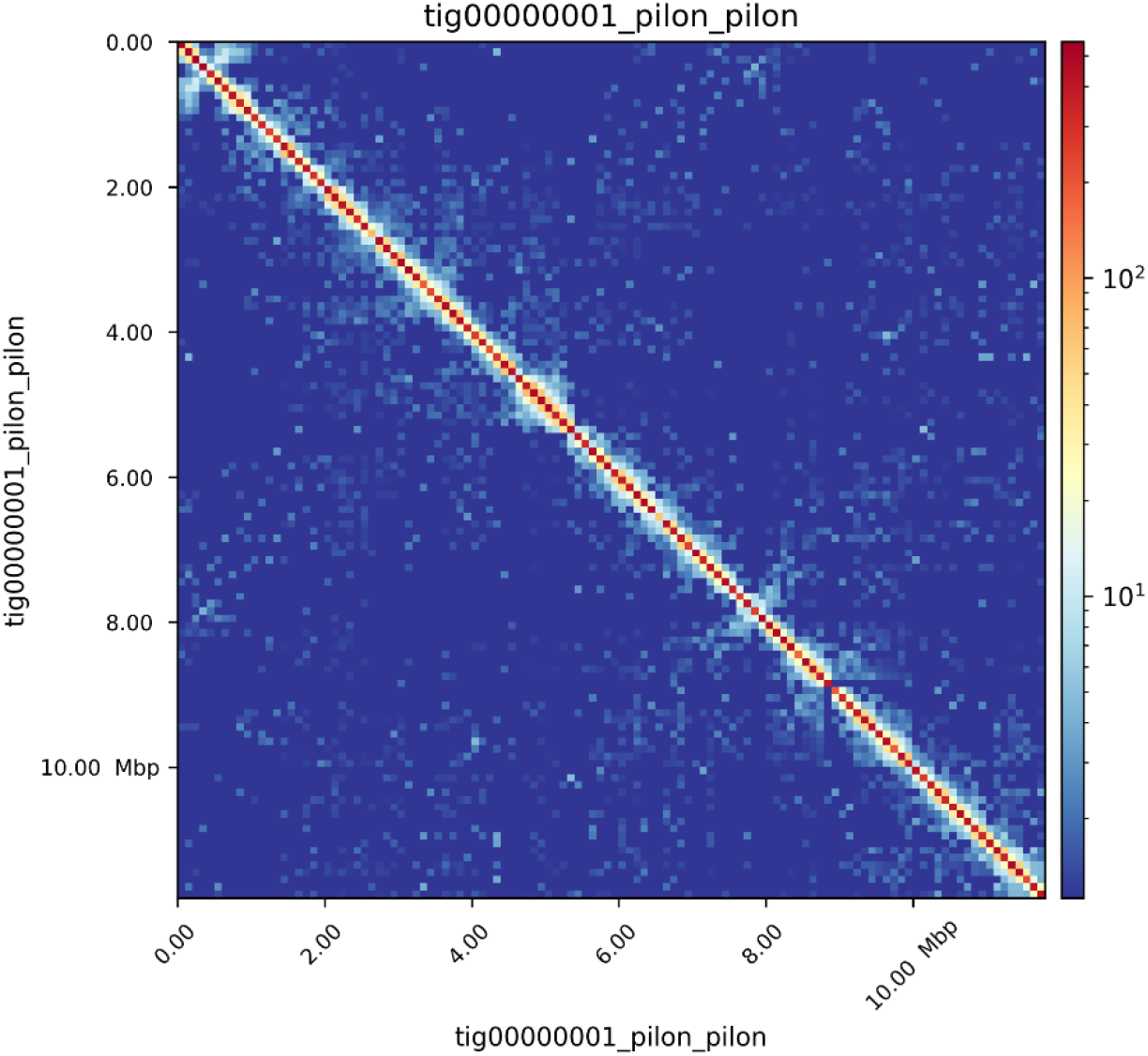
Hi-C contact map of a chimeric contig in the Nanopore assembly that was broken prior to phasing and scaffolding. **(A)** In the Nanopore assembly, contig tig00000001 has two centromeric regions visible at ~0.5 Mb and ~7.8 Mb. The chimeric breakpoint was chosen at 5.35 Mb based on visual inspection of long-read alignments.

**Supplementary Figure S2:**
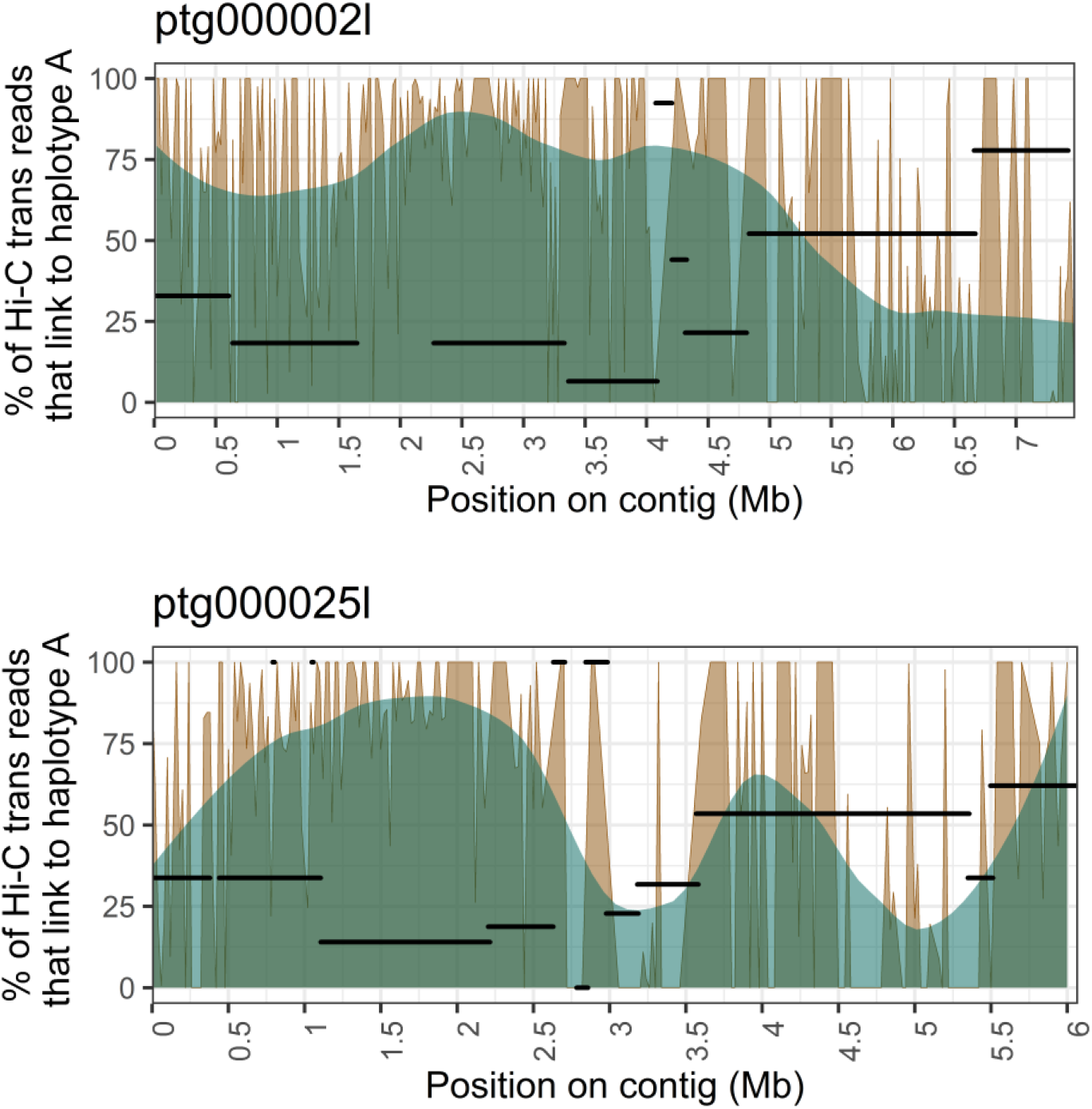
Contigs ptg000002l and ptg000025l from the HiFi-hifiasm assembly and its associated haplotigs (black segments). The % of Hi-C *trans*-contacts that link to haplotype A (with an associated smoothing line) are shown. Haplotigs are shown at the *y* coordinate that corresponds to their % of Hi-C *trans-*contacts to haplotype A. If a haplotig has no or only few Hi-C *trans*-contacts, it is shown at *y* = 100. Contig ptg000002l appears to switch phase at ~5.5 Mb, which does not clearly overlap with the corresponding haplotig alignment start and end points. Similarly, the phase switch point at ~4.5 Mb in contig ptg000025l does not clearly overlap with the corresponding haplotig alignment start and end points.

**Supplementary Figure S3:**
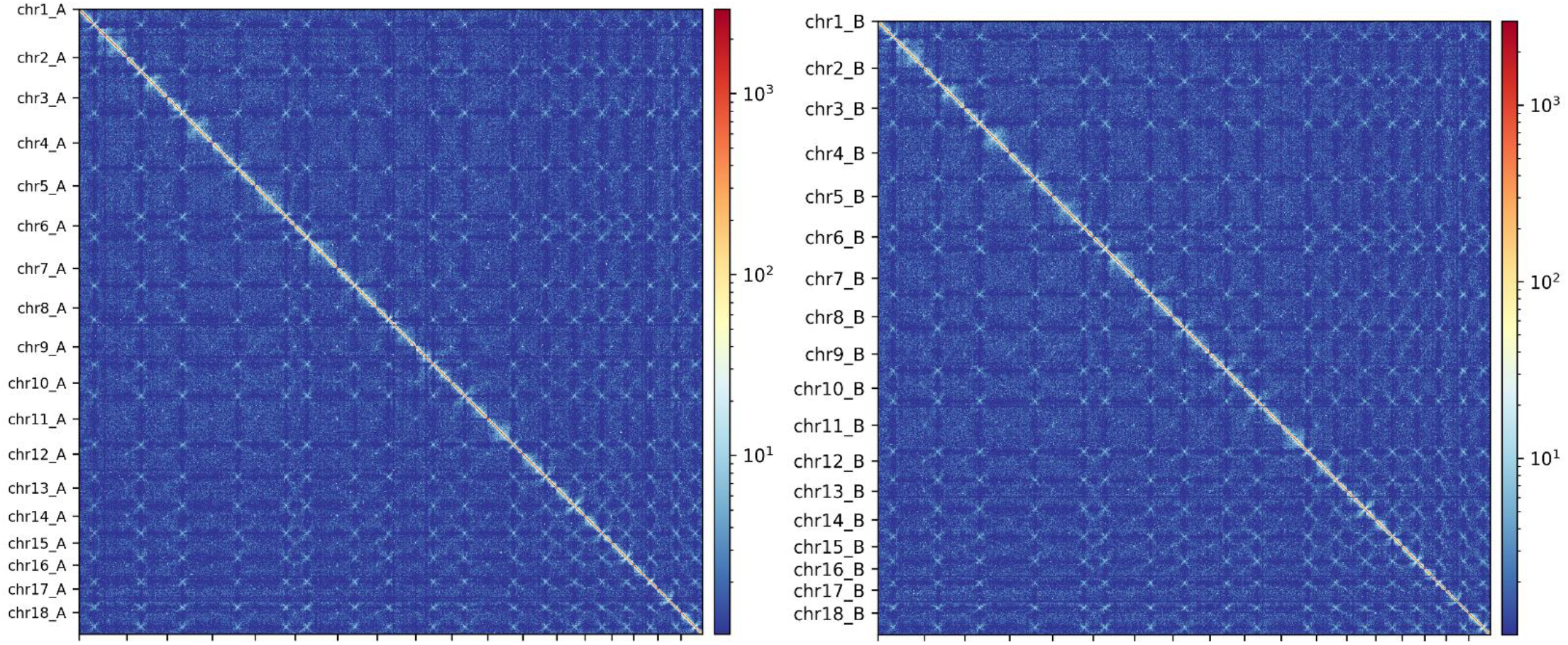
Hi-C contact maps of the two chromosome haplotypes in the HiFi assembly. The centromeres are visible as distinct cross-shapes in the whole haplotype Hi-C contact maps.

